# Individual Molecular Motors use Low Forces to bypass Roadblocks during Collective Cargo Transport

**DOI:** 10.1101/2021.01.12.425491

**Authors:** Saurabh Shukla, Alice Troitskaia, Nikhila Swarna, Barun Kumar Maity, Marco Tjioe, Carol S Bookwalter, Lynn Chrin, Christopher L Berger, Kathleen M Trybus, Yann R Chemla, Paul R Selvin

**Affiliations:** Department of Chemical and Biomolecular Engineering, University of Illinois at Urbana-Champaign, Urbana, Illinois; Center for the Physics of Living Cells, University of Illinois at Urbana-Champaign, Urbana, Illinois; Center for Biophysics and Quantitative Biology, University of Illinois at Urbana-Champaign, Urbana, Illinois; Department of Physics, University of Illinois at Urbana-Champaign, Urbana, Illinois; Department of Molecular Physiology and Biophysics, University of Vermont, Burlington, VT

## Abstract

A cargo encounters many obstacles during its transport by molecular motors as it moves throughout the cell. Multiple motors on the cargo exert forces to steer the cargo to its destination. Measuring these forces is essential for understanding intracellular transport. Using kinesin as an example, we measured the force exerted by multiple stationary kinesins *in vitro*, driving a common microtubule. We find that individual kinesins generally exert less than a piconewton (pN) of force, even while bypassing obstacles, whether these are artificially placed 20-100 nm particles or tau, a Microtubule Associated Protein. We demonstrate that when a kinesin encounters an obstacle, the kinesin either becomes dislodged and then re-engages or switches protofilaments while the other kinesins continue to apply their (sub-)pN forces. By designing a high-throughput assay involving nanometer-resolved multicolor-fluorescence and a force-sensor able to measure picoNewtons of force, our technique is expected to be generally useful for many different types of molecular motors.

## Introduction

Molecular motors produce mechanical forces to transport cargo inside the cell and are vital for many cellular processes such as cell division, cell growth, and distribution of cellular resources^1,2^. Typically, multiple molecular motors are attached to a single cargo^3,4^ and dynamically coordinate their forces with each other to move the cargo to its destination^5,6^, oftentimes overcoming cellular roadblocks, such as other filaments or other proteins (such as tau, a microtubule-associated protein (MAP)^7,8^). Using *in vitro* fluorescence assays, it has been shown that single kinesin motors are inhibited by artificial and physiological roadblocks but can bypass the roadblock when they work in teams^6,9^. *In vitro* single-molecule fluorescence assays can quantify the position, run lengths, and velocities of the cargo but cannot determine the individual properties of each motor—particularly the force—as it participates in transport. Quantitative measurement of motor forces during collective cargo transport is essential for understanding motor motility and the mechanism of bypassing cellular roadblocks.

Atomic force microscopy, magnetic tweezers, and optical traps are powerful techniques to quantify forces of biomolecules at the single-molecule level. Out of these, the optical trap assay has been the most widely used technique to study the responses of molecular motors to applied forces^10–12^ and to measure motor stall forces^13–15^. In this assay, a microsphere attached to the molecule of interest is captured one at a time, resulting in low-throughput data collection. Optical traps can characterize the collective force-response of multiple motors that are attached to a single microsphere but cannot quantify forces by individual motors in this configuration^13,16^. Additionally, assessing the impact of diverse experimental configurations such as the presence of roadblocks on individual motor force generation is also difficult with existing force sensing techniques. A high-throughput force sensor capable of resolving the forces generated by individual molecular motors during collective transport will enable new insights and an in-depth understanding of cellular cargo transport.

Recently, several single-molecule force sensing techniques have been developed with DNA-based designs^17^. Some examples include quantifying forces of antigen-antibody interactions^18^, receptorligand interactions^19,20^, nucleic acid systems such as Holliday junctions^21^, nucleosome complexes^22^, and protein-DNA interactions^23^. Recently, a nanospring made out of DNA origami was used to measure myosin forces^24^. Here, we develop a Force Sensing by Individual Motors (FSIM) assay to measure forces of kinesins collectively moving a common microtubule, as well as the displacement and velocity of the common cargo. Our assay is based on quantifying the displacement of quantum dot labeled kinesin using fluorescence and calculating the force by measuring the extension of a denatured single stranded DNA molecule. Force data of tens to hundreds of kinesins can be measured in a single microscope movie with our assay. The developed force-sensing assay provides high-throughput measurements of forces and is expected to be useful with other types of molecular motors such as myosin and dynein. Our assay enables force resolution at the single motor level during collective cargo transport, a task that is difficult to achieve with conventional assays.

## Results

### Design of FSIM assay

The FSIM assay was closely based on our previous study, where we measured the displacement of multiple kinesins held down independently to a coverslip as they simultaneously moved a common microtubule cargo^6^ (see Fig. 1a). The displacement of individual kinesins was measured by Fluorescence Imaging with One Nanometer Accuracy (FIONA)^25^ of a quantum dot attached to the end of kinesin which was placed on the coverslip via a flexible double-stranded DNA (ds-DNA). To measure the force exerted by the kinesins, we needed a reliable “spring” of known elastic behavior to convert the displacement, x, into a force F. We found that a ds-DNA was not an effective force-sensing molecule at high forces; small errors in measured extension (determined by imaging) translate to large errors in force calculation (Fig. S1).

**Figure 1.**
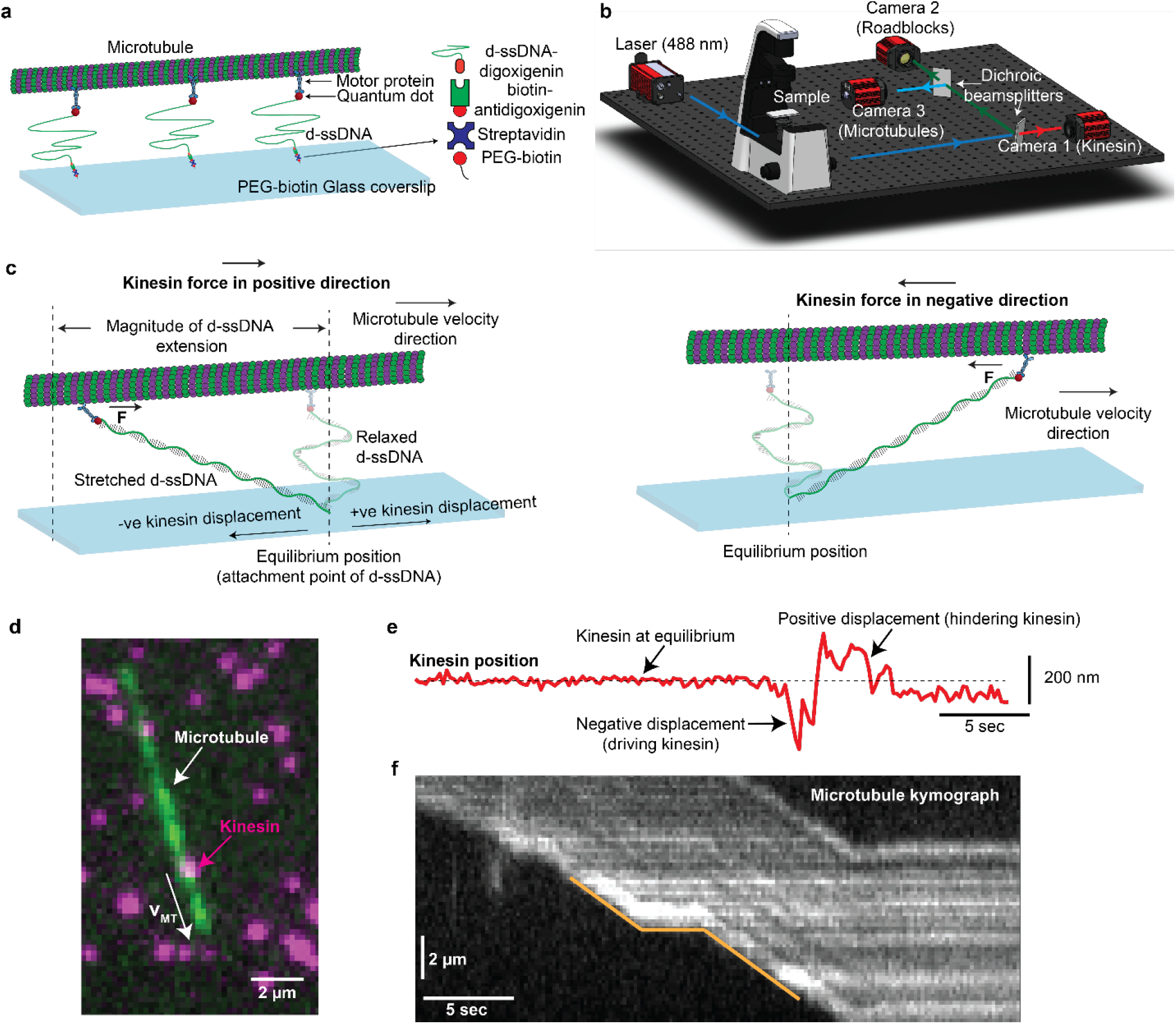
Design of the FSIM assay. **(a)** Configuration of our FSIM assay where multiple kinesins can simultaneously transport a common microtubule. Kinesins are attached to the surface using a d-ssDNA linker and are placed sufficiently away from each other to be tracked individually. **(b)** Schematic of the microscope setup where the signal from kinesins, microtubules, and the roadblocks can be captured in three separate cameras. We label kinesins, microtubule, and the roadblocks with spectrally different fluorescent tags. **(c)** Schematic depicts the force calculation procedure for individual kinesins. By convention, displacements in the direction of the microtubule velocity are assumed to be positive displacements. Kinesin exerts force in the forward direction (direction of the microtubule velocity) when it has negative displacement (left schematic). d-ssDNA extends when kinesin starts producing force after coming into contact with the microtubule. Kinesin force is calculated from the d-ssDNA extension as determined from the fluorescent signal of the quantum dot. The right schematic shows the kinesin molecule exerting force in the backward direction. **(d)** Microscopic image of a microtubule (green) driven by two kinesins (magenta). The white arrow represents the direction of microtubule velocity. **(e)** The position of a kinesin molecule along the microtubule axis is shown. The black arrows depict the equilibrium position, positive displacements, and negative displacements. Forces of individual kinesins are derived from their displacements around the equilibrium position. **(f)** Kymograph of the microtubule being driven by surface kinesins. The velocity of the microtubule is calculated by tracing the kymographs (see yellow tracing lines).

In contrast, we found that denatured single-stranded DNA (d-ssDNA) worked well in the range of 0-15 pN (Fig. 1a). Denaturation of the ssDNA with glyoxal was necessary to eliminate any secondary structures which can cause instantaneous discontinuities in the force-extension behavior when pulling^26^. By labeling the kinesins on the surface reasonably sparsely, the kinesin motors were placed sufficiently far-away from each other such that no two kinesins were in the same diffraction-limited spot. We optimized our experimental protocol such that one quantum dot was attached to one kinesin and one d-ssDNA molecule to ensure the accuracy of our force measurement (see methods). We used our MiCA purification method developed in-house^27^ and proper controls (Fig. S2) to minimize non-specific binding of free quantum dots and free kinesins on the surface. We labeled kinesins, microtubules, and the roadblocks (introduced later in the paper) with QD705, Hilyte 488, and QD605, respectively. Since these labeled tags were spectrally different, we could simultaneously image kinesins, roadblocks, and microtubules in three different cameras (Fig. 1b). Owing to the high photostability of quantum dots and the high labeling density of microtubules, we could image for several minutes (>5 minutes) to collect force data of hundreds of kinesins per movie.

To demonstrate the functionality of our force sensor, we observed microtubule transport by d-ssDNA-linked kinesin motors. Due to the stretchability of d-ssDNA, kinesins were not rigidly immobilized on the surface. They could be displaced from the attachment point of the d-ssDNA (defined as the equilibrium point of the kinesin). A kinesin could translocate in either direction (towards or opposite to the microtubule velocity) with respect to its equilibrium position and assume a driving or hindering state (Fig. 1c) during cargo transport. In the driving state, kinesin translocated in the opposite direction of the microtubule velocity (negative displacement) and applied a positive force for propelling the microtubule in the forward direction (Fig. 1c left schematic). In the hindering state, kinesin translocated in the direction of the microtubule velocity (positive displacement) and applied a negative force (in the opposite direction of microtubule velocity; Fig. 1c right schematic and Fig. S3).

Fig. 1d shows a microscope image of kinesins held down via a d-ssDNA to the coverslip and a microtubule. We tracked the positions of individual kinesins with time. Fig. 1e shows the position trace of one of the kinesins where its equilibrium, driving, and hindering position can be seen. Kinesins remained at their equilibrium point in the absence of microtubules (dashed horizontal line). When a microtubule approached the kinesins, the kinesins were displaced from their equilibrium point as they walked on the microtubule, thereby stretching the d-ssDNA. The displacement of each kinesin from the equilibrium position caused a change in the extension of d-ssDNA. Hence, the force exerted by each kinesin could be estimated based on the d-ssDNA extension. We note that the vertical distance between kinesin and the surface was expected to be small and contributed minimally to the extension of d-ssDNA and the force (see supplementary text appendix 1). Therefore, the displacement of each kinesin from its equilibrium position was assumed to be equal to the d-ssDNA extension. We also tracked the microtubule position in a different camera and calculated the microtubule’s velocity by tracing its kymograph (Fig. 1f).

### Synthesis and force-extension characterization of d-ssDNA

For the experimental realization of our assay, we needed a dual-functionalized single-stranded DNA. Using a series of genetic engineering approaches, we synthesized monodisperse dsDNA with one strand functionalized with biotin at one end and with digoxigenin at the other end (Fig. 2a and Methods). Subsequently, we denatured the dsDNA with glyoxal and obtained our desired 1180 bp long d-ssDNA product. Glyoxal covalently binds to the DNA base pairs and prevents secondary structures, making the d-ssDNA a reliable construct for force sensing applications^26,28^. Secondary structures introduce conformational variability amongst ssDNA molecules that translate to inconsistent force measurement. The length of d-ssDNA (1180 bp with ~500 nm contour length) was chosen such that each kinesin could move sufficiently around its equilibrium position without interfering with other kinesins.

**Figure 2.**
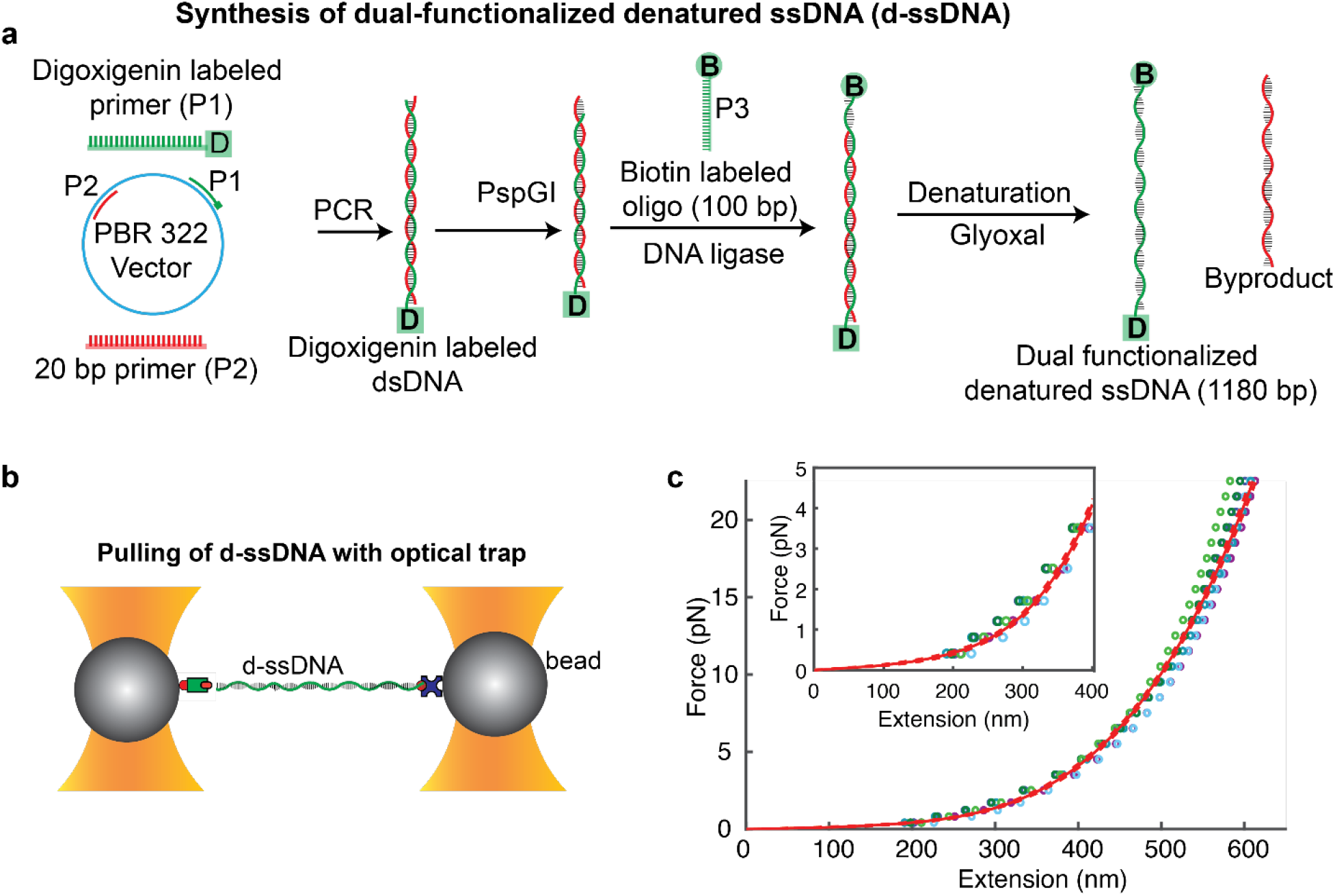
Synthesis and characterization of d-ssDNA. **(a)** Synthesis of a mono-dispersed d-ssDNA functionalized with biotin at one end and digoxigenin at another end. Glyoxal is used to synthesize denatured ssDNA (d-ssDNA) and to ensure minimal secondary structure formation. **(b)** Schematic of the optical trap assay for experimentally determining the force-extension curve of the d-ssDNA. **(c)** Force-extension curves were obtained from the optical trap assay. Force-extension curves are fitted with an analytical model (dashed lines very close to fit are 95% confidence intervals for fit). This reference FEC is used to determine the forces of individual kinesins from their respective displacements. A zoomed in view of the model at low forces is also shown.

We characterized the force-extension characteristics of our d-ssDNA molecule using an optical trap assay (Fig. 2b). We pulled on individual d-ssDNA molecules using dual-trap optical tweezers^29,30^ where a d-ssDNA-coated streptavidin microsphere was captured in one trap and a microsphere coated with anti-digoxigenin antibody in the other. We obtained force-extension curves (FECs) of d-ssDNA molecules in the same imaging buffer used for fluorescence measurements. The majority of FECs (Fig. 2c and S4) fit well to an analytical expression based on previously published data^28,31^ (see methods). We used the obtained analytical expression as our reference model for calculating the forces of kinesin motors in our force-sensing assay (Fig. 2c). Unlike the FEC of dsDNA, the d-ssDNA FEC has a moderate slope that makes our d-ssDNA construct a more sensitive and reliable force sensing molecule for molecular motors in the relevant force range of 0-10 pN (Fig. 2c and S1).

### Forces by individual kinesin motors during multi-motor transport

An example of a microtubule (green) transport by three kinesin motors (magenta, enclosed in white circles) is shown in Fig. 3a. As soon as the microtubule became attached to the kinesins, the kinesins began walking on the microtubule and are moved from their equilibrium positions. We plotted the microtubule velocity (Fig. 3b top panel) and the displacements of each of the three kinesins with time (Fig. 3b middle panels). Using kinesin displacements and reference FEC, we estimated the force exerted by each kinesin on the microtubule during collective transport (Fig. 3b bottom panels). We observed that individual kinesins’ forces varied dynamically with time. For example, kinesin #1 and #3 applied positive forces intermittently, while kinesin #2 remained at its equilibrium position under the microtubule for the entire transport duration but exerted negligible forces.

**Figure 3.**
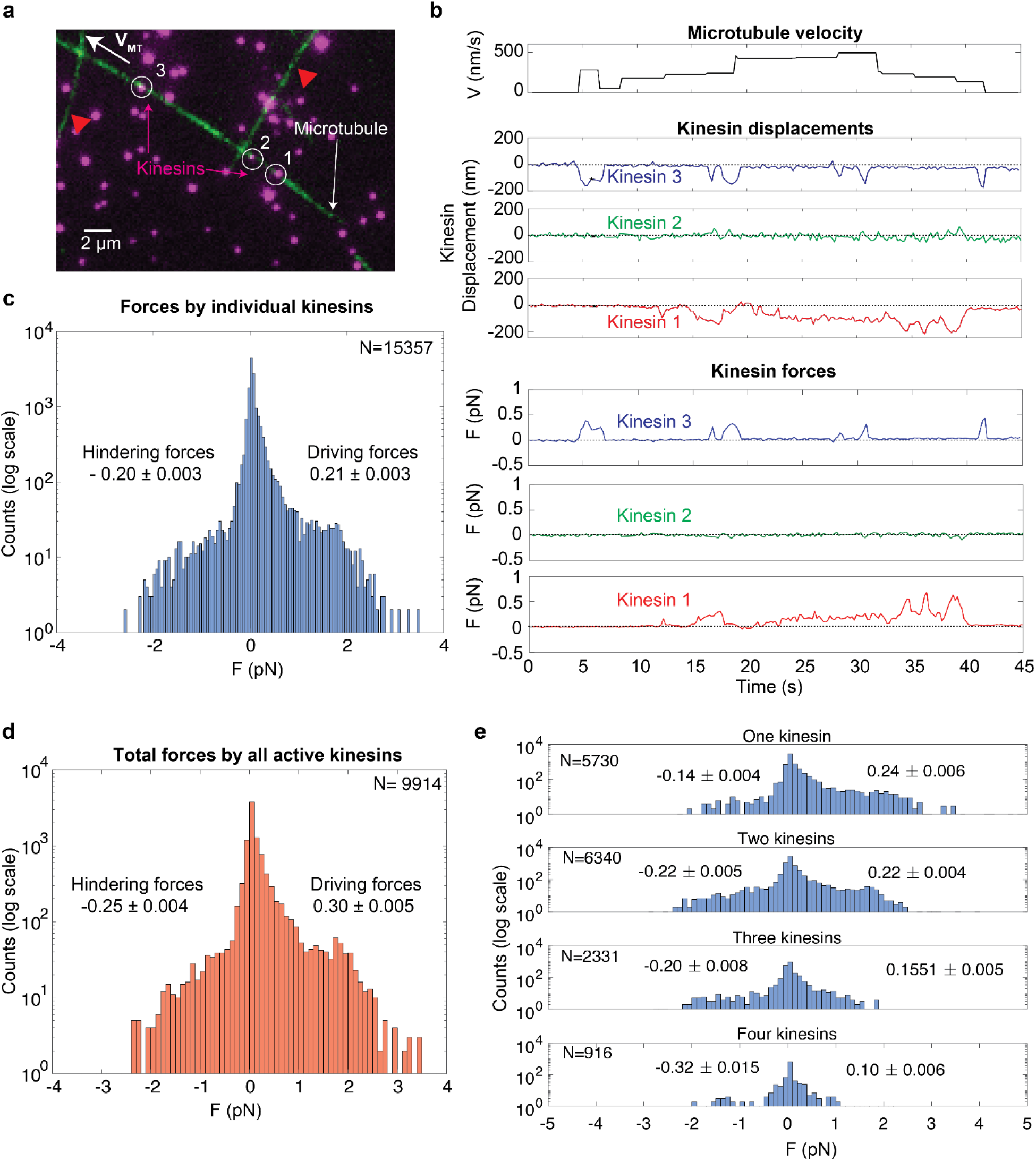
Forces by individual kinesin motors. **(a)** Force dynamics of individual kinesin motors in a case where three kinesins (magenta, enclosed with white circles) are simultaneously transporting a common microtubule (green). The white arrow (VMT) shows the direction of microtubule velocity. Red triangles indicate two other microtubules that interact with the microtubule transported by three kinesins. **(b)** Microtubule velocity, kinesin displacements, and individual forces of kinesins. **(c)** Histogram of force produced by individual kinesin motors during cargo transport. Forces are recorded when a kinesin is transporting a microtubule. Kinesins produce an average of ~0.21 pN force in the forward direction and ~0.20 pN of force in the backward direction (mean ± s.e.m.). **(d)** Total force by all the kinesins that are actively exerting force at a particular time point on the microtubule is plotted. The average total force on the microtubule is 0.30 pN in the forward and 0.25 pN in the backward direction. **(e)** Average forces exerted by individual kinesin motors with the varying number of total kinesins. The average forward force per kinesin decreases as the number of kinesins increases. The average backward force increases as the number of kinesins increases.

To make sense of the forces by individual kinesins, we carried out a control, measuring the forces by kinesins at their equilibrium position (Fig. S5). When there is no microtubule attached to kinesins, kinesins remain at their equilibrium position, and the forces by kinesins are close to zero. We also observed microtubule transported by a single kinesin (Fig. S6). Forces by one attached kinesin remain very low. In fact, this makes sense: For transporting a microtubule, one kinesin needs to produce a force comparable to the drag force on the microtubule, which is expected to be < 0.1 pN (see supplementary text appendix 2)^32^. In the case of Fig. 3a, the microtubule moved by three kinesins interacted with two other microtubules, which prompted kinesins to increase their forces for the continuation of the transport, as shown in Fig. 3a (red triangles) and Supplementary Movie 1. At other times, e.g., from t = 7 to 12 sec, the microtubule is moving with no appreciable load, i.e., the forces exerted by kinesins are undetectably small.

To delve into the statistics of force exerted by kinesin motors during collective transport, we analyzed 82 instances (82 independent microtubules) where a microtubule was being transported by multiple kinesins and obtained thousands of kinesin force data points (N=15357, N represents the number of the force data points with the frame rate of 0.2 sec; see also Fig. S7-8 for discussion of other cases). We plotted the histograms of forces produced by individual kinesin motors (Fig. 3c). Forces were only recorded when a microtubule was attached to the kinesin, and the kinesin was transporting (either driving or hindering) the microtubule. Kinesins exerted forces in the forward direction most of the time— ~76% of the force data points were in the direction of microtubule velocity— and ~24% of forces were in the negative direction and hindered the transport of the microtubule. Kinesins exerted an average of ~0.21 pN force during the collective transport of microtubules in the positive direction and an average of ~0.20 pN of force in the negative direction (Fig. 3c). We also calculated the total forces on the microtubule by adding the forces of all the kinesins that either drove (+) or hindered (-) the microtubule motion at a given point of time. The average total force exerted by the kinesis was ~0.30 pN in the forward direction and was marginally higher than the average individual kinesin force of ~0.21 pN (Fig. 3d). Although individual kinesins exert forces up to ~3 pN occasionally (see Fig. S9), they need to produce much smaller forces on an average for transporting the cargo than its stall force of 5-7 pN^33–35^.

Our assay also allowed us to quantify the effect of the number of kinesins on individual kinesin forces. We segregated the cases where a different number of kinesins were actively transporting the cargo and estimated their average force. Fig. 3e shows the individual kinesin forces when one, two, three, and four kinesins exert forces on the microtubule. As the total number of kinesins increases on the microtubule, the average force per kinesin in the positive direction decreases, indicating that driving kinesins share the load amongst themselves during the collective cargo transport. Interestingly, the average negative force by kinesins increases with the increasing number of total kinesins indicating that hindering kinesins do not share the load among themselves. As discussed earlier, the number of driving kinesins is almost three-times that of hindering kinesins. As the total number of kinesins on the cargo increases, the hindering kinesins must balance the forces by a higher number of driving kinesins, and therefore need to exert higher forces.

### Kinesin forces in the presence of quantum dot roadblocks

In cells, which tend to be extremely crowded with proteins (~300 mg/ml)^36,37^, molecular motors need to overcome many roadblocks to reach their destination, including other cytoskeletal roadways and various microtubule-associated proteins (MAPs). To mimic cellular roadblocks, we attached fluorescent quantum dots (QD605, ~20-25 nm in size) to the microtubules using streptavidin-biotin linkages and used these QD-decorated microtubules in our force sensor assay. We mixed 50 nanomolar (nM) of QD-roadblocks with an equal volume of microtubules (5 μM of polymerized microtubules), which corresponded to ~1.6 QDs per micron length of the microtubule (See methods). We could track the dynamics of kinesins, roadblocks, and microtubule simultaneously using a three-camera experimental setup (Fig. 1b).

To quantify the forces exerted by kinesins in the presence of the roadblocks, we analyzed the movement of kinesins and QD-decorated microtubules. We observed that as roadblocks were introduced in the system, the motion of the microtubule became less smooth, and there were more occurrences of stopping and bending of microtubules. We illustrate one of the cases where four kinesins transport a microtubule in Fig. 4a. Initially, the microtubule attached to kinesins and started moving and we can see the microtubule and the roadblocks approaching the kinesins (Fig. 4a left, t ~ 6 sec). At t ~ 8 sec, one kinesin became stuck at the roadblock (Fig. 4a middle). Eventually, at t = 12 sec, the kinesins overcame the roadblock, and the microtubule started moving again (Fig. 4a right; snapshot shown at t = 16 sec).

**Figure 4.**
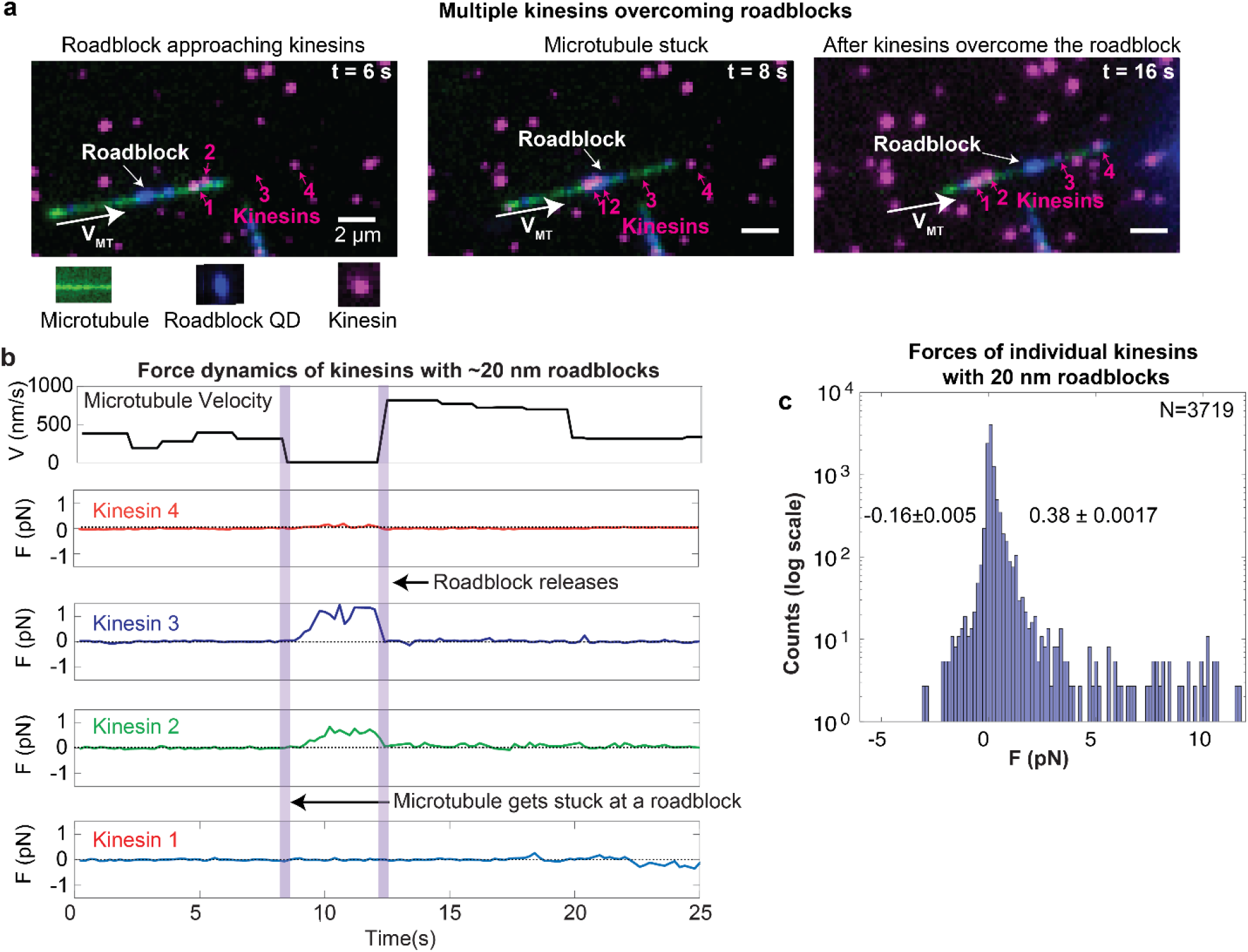
Force dynamics of kinesins in the presence of ~20-nm quantum dots as roadblocks. **(a)** Microscope images of an instance at different time points where four kinesins overcome a roadblock. Microtubule, kinesins, and roadblocks are shown in green, magenta, and blue colors, respectively. The white arrow shows the microtubule velocity direction. One of the kinesins (kinesin #2) becomes stuck on the roadblock on the microtubule at t = 8 s (middle image). The rightmost image illustrates when kinesins have overcome the roadblock, and the microtubule starts moving again. The fluorescence intensity of kinesin #3 is dim in the first two images as it is out of focus. **(b)** Forces by four kinesins are plotted when they are simultaneously moving a common microtubule. Microtubule velocity is also plotted. The vertical purple rectangles depict the time points when one kinesin becomes stuck at the roadblocks (t ~ 8 s) and when the roadblocks get released (t ~ 13 s). Microtubule velocity resumes after kinesin #1, #2 and #3 apply forces between t~8 sec and t~13 sec. **(c)** Force histogram of kinesins in the presence of roadblocks. Kinesins exert ~0.38 pN and ~0.16 pN force on an average in the forward and backward direction, respectively, in the presence of the roadblocks (mean ± s.e.m.).

We plotted the forces exerted by each kinesin and the microtubule velocity with time (Fig. 4b). One of the kinesins (kinesin #2) became stuck at the roadblock at t~8 sec (vertical purple rectangle in Fig. 4b). At this point, kinesin #2, #3, and #4 exerted higher forces, and kinesin #1 did not increase its force. The stuck kinesin bypassed the roadblock, and the microtubule started moving again at time = 12 sec. After kinesin #2, #3, and #4 applied forces, the stuck kinesin (#2) either changed protofilaments or dislodged from the microtubule and reattached to the microtubule for bypassing the roadblock. After bypassing the roadblock, the forces of all the kinesins decreased as the kinesins needed to apply only low forces, those comparable to the drag forces on the microtubule. Forces of the three kinesins remained much below the stall force even when a kinesin got stuck at the roadblock. This instance demonstrates that multiple kinesins can simultaneously increase their forces marginally, overcoming the roadblocks (See supplementary movie 2). We discuss more cases where kinesins dynamically change their force in the presence of roadblocks in the supplementary text (Fig. S10-12).

We analyzed 40 instances (40 independent microtubules) of the roadblock-loaded microtubule transport by multiple kinesins to obtain the aggregate force characteristics (Fig. 4c). We plotted the force exerted by individual kinesin motors in the presence of roadblocks. Forces were only recorded when a kinesin interacted with the microtubule. We found that the average force exerted by kinesin motors marginally increases when the quantum dot roadblocks are introduced in the assay. In addition, with roadblocks, kinesin occasionally reached forces comparable to its stall force (1.4% force data points were >5 pN) but primarily remained in the sub-pN regime (Fig. 4c). Kinesins produced an average of ~0.38 pN in the positive direction and ~0.16 pN in the negative direction with QD605 as roadblocks. These results show that kinesins primarily exert less than 1 pN forces even in the presence of the roadblocks and rarely, but occasionally, increase their forces to their stall forces. (Fig. S9).

### Kinesin forces with larger microspheres

Next, we asked if the force exerted by kinesin increases if we increase the size of the roadblocks. Therefore, we used 100-nm microspheres (compared to the ~20 nm QD) as roadblocks. One such case where kinesin became stuck at the roadblock and eventually overcame (it is described in Fig. 5a, b). In this case, three kinesins transported a single microtubule. When kinesin #2 became stuck at the roadblock at t = 2.6 s, two kinesins (#2 and #3) increased their forces but remained less than 1 pN to resume the microtubule motion (See supplementary movie 3). We analyzed 38 such cases and plotted the force histogram (Fig. 5c). Kinesins exerted an average force (per kinesin) of ~0.43 pN in the forward direction and ~0.16 pN in the backward direction, which is comparable to the forces in the presence of QD605 as roadblocks (Fig. 4c). Therefore, even in the presence of bigger roadblocks, the average forces exerted by kinesins while interacting with roadblocks remained below 1 pN. Our results show that the kinesins did not need to increase their forces to overcome bigger roadblocks. We speculate that kinesins overcome the roadblocks by detaching from the microtubule and immediately reattaching after bypassing the roadblock and/or switching the protofilaments while keeping their forces low.

**Figure 5.**
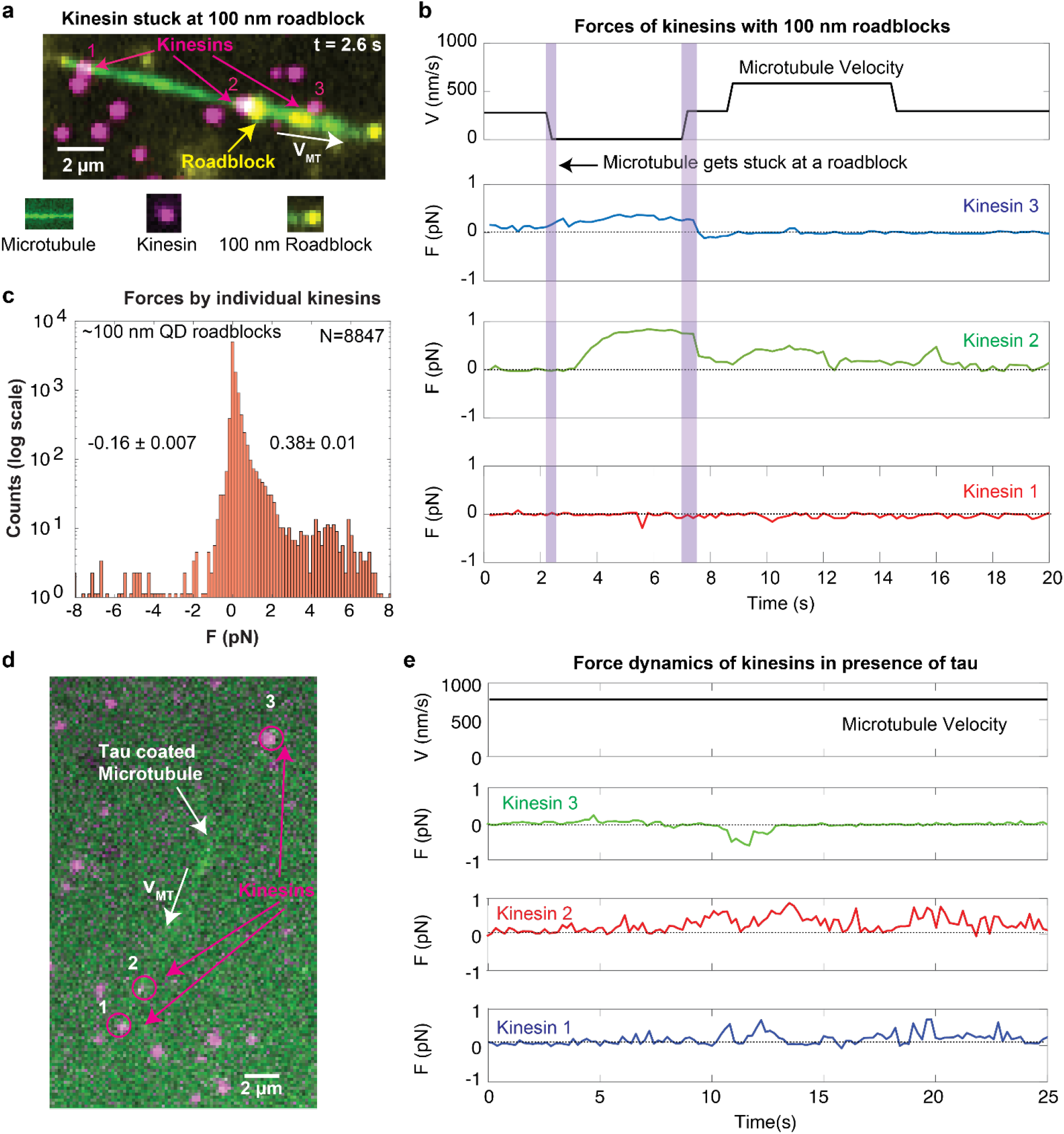
Kinesin forces in the presence of 100-nm roadblocks and tau. **(a)** Microscope image where kinesins overcome a 100-nm roadblock. The microtubule (green) is driven by three kinesins (magenta). The roadblocks attached to the microtubules are yellow. Kinesin #2 becomes stuck at the roadblock. **(b)** Microtubule velocity (top) is plotted with kinesins’ forces for the case shown in Fig. 5a. At t = 2.6 s, kinesin #2 becomes stuck at the roadblock, and the microtubule stops. Kinesin #2 and #3 increase their forces and overcome the roadblock at t = 7.5 s. Purple rectangles indicate the regions when the kinesin becomes stuck at the roadblock and are released. Kinesin #1 does not increase its force and does not play a role in overcoming the roadblock. Forces of kinesins remain below 1 pN for overcoming the roadblock. **(c)** We record the forces exerted by kinesins transporting the roadblock (~100 nm QDs) loaded microtubules (38 independent microtubules). In the presence of 100 nm roadblocks, the mean forces of kinesins remain 0.38 pN and 0.16 pN in the forward and backward direction, respectively (mean ± s.e.m.). **(d)** Microscope image of an example where three kinesins (magenta) are transporting a tau coated microtubule (green). A high concentration of tau (100 nM) is used in the assay, and therefore, the entire surface of the microtubule is coated with tau. The microtubule velocity (vMT) direction is shown with an arrow. **(e)** Microtubule velocity, along with the individual kinesin forces, is shown. All three kinesins exert forces less than 1 pN for transporting the tau coated microtubule.

### Multiple kinesins can transport the cargo in the presence of tau

We also performed the FSIM assay with the physiological roadblock tau. We incubated microtubules with excess Alexa532-labeled 3RS tau (100 nM in the imaging buffer, see methods) such that tau completely coated the microtubules. 3RS tau is the shortest of the six tau isoforms expressed in the human brain^38^. Fig. 5d and 5e show an example of three kinesins (#1,2,3) transporting a tau-coated microtubule and the microtubule velocity and the forces exerted by kinesin #1, 2, and 3 (see supplementary movie 4). Forces primarily remain below 1 pN, and the microtubule has an average velocity of ~760 nm/sec. Kinesin #1 and #2 apply forces mostly in the positive direction, while kinesin #3 exerts a negative force for a short time (at t=11 s). One more example of tau coated microtubule transport by four kinesins is shown in Fig. S13. Despite the previous understanding that tau inhibits kinesin-based transport^39–43^, our results show that multiple kinesins can transport the microtubule in the presence of tau. With our assay, we can also measure forces exerted by individual kinesins in real-time in the presence of tau. These experiments show that the FSIM assay is versatile and can be used in various artificial and physiologically relevant roadblock configurations.

## Discussion

In this work, we have leveraged the power of fluorescence microscopy to develop a d-ssDNA-based Force Sensing of Individual Motors (FSIM) assay. Our assay can visualize individual motors and measure their forces during multi-motor transport of a common microtubule cargo. We performed optical trap assay measurements to characterize the force-extension properties of the d-ssDNA molecule that acts as a force sensing molecule in the FSIM assay. Our FSIM assay adds a high-throughput and single-motor resolution method to the repertoire of available force spectrometry techniques. Our assay is versatile and can be used to study the force dynamics of motors in diverse configurations, such as the presence of a variety of roadblocks. We have shown the functionality of the FSIM assay with two sizes of artificial roadblocks (~20-nm quantum dots and 100-nm microspheres) and physiologically relevant microtubule-associated protein tau. With our assay, we have demonstrated that kinesin motors modulate their forces to overcome roadblocks. Kinesin produces enough force to overcome drag on the cargo while occasionally increasing its forces. We have shown that kinesins keep their forces primarily below one pN and can still overcome the roadblocks. We propose that kinesins’ mechanism of roadblock evasion is predominantly detachment from microtubule or switching of protofilaments. By operating with multiple kinesins driving a cargo, the cargo can be moved when tau or other molecules would make directed motion difficult. Furthermore, our FSIM assay is expected to be useful for understanding the motion of opposite-directed and different motors.

## Methods

### Synthesis of glyoxal denatured single-stranded DNA (d-ssDNA)

The process of d-ssDNA synthesis is shown in Fig. 2a. pBR322 plasmid (NEB, catalog #N3033S) was used as a vector for PCR amplification of 1116 bp long dsDNA using primers P1 and P2 (sequences of all primers are given in Table S1). Primer P1 had a digoxigenin modification at its 5’ end. All primers were ordered from IDT. The PCR product was digested with PspGI restriction enzyme (NEB, catalog #R0611S) to obtain a 1080 bp long DNA with 5 bp overhang and was subsequently dephosphorylated with Antarctic phosphatase (NEB, catalog #M0289S). The obtained product was PCR purified using Qiagen PCR purification kit, and concentration was measured using Nanodrop One (ThermoFisher). The obtained DNA product (1080 bp) was annealed with primer P3 (100 bp long) at a 1:3 DNA to primer molar ratio and ligated using T4 DNA ligase (NEB, catalog #M0202S). Primer P3 had biotin modification at its 3’ end. After the ligation, we obtained a dsDNA product that had one strand functionalized with digoxigenin at one end and biotin at another end. The length of this strand was 1180 bp.

Next, we denatured the DNA with glyoxal^26^ to get the final denatured single-stranded DNA, d-ssDNA. First, glyoxal was deionized using mixed bed resin (AG 501-X8, BioRad, Hercules, CA, USA). 10 ml of glyoxal was treated with 2g of mixed bed resin and was stirred for 2 hours. The deionization process was repeated three times.

Glyoxal treatment of 1180 bp long dsDNA was carried out in TE buffer (ThermoFisher) with a reaction volume of 20 ul. Reaction volume contained 50% v/v DMSO and 1 M glyoxal. Glyoxal treatment was carried out at 50°C for 1 hour in a PCR thermocycler (Eppendorf). The final product was run on 1% agarose gel in TBE buffer for 1 hour at 100V. d-ssDNA on the gel was stained with SYBR gold stain and was gel purified with a QIAquick gel extraction kit (Qiagen) (Fig. S14). The concentration of d-ssDNA was measured with NanoDrop One (ThermoFisher).

**Table S1.**
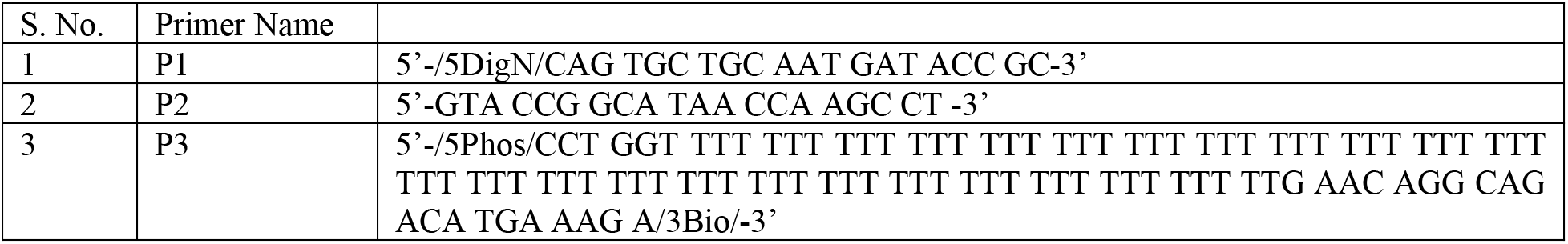

### Optical Trapping Assay

The stretching behavior of glyoxalated single-stranded DNA (d-ssDNA) in the same buffer conditions as the microtubule assay was characterized by pulling on individual molecules using dual-trap optical tweezers^29,30^.

Glyoxalated single-stranded DNA molecules identical to those used in the microtubule assay (i.e., 1180-nt long, functionalized with 3’ biotin and 5’ digoxigenin) were synthesized as described earlier above. For optical trapping experiments, d-ssDNA was diluted to 0.14 nM, and 1 to 1.5 μL of the d-ssDNA solution was incubated for an hour at room temperature with 5 μL of 0.2% w/v streptavidin-coated microspheres (Spherotech). Beads were diluted in approximately 300 μL of buffer (100 mM Tris, 20 mM NaCl, 3 mM MgCl2, pH 7.6) for delivery to the optical traps through bead channels in a custom flow chamber^44^.

The trapping channel of the flow chamber contained buffer consisting of: 91% DmB-BSA (30 mM HEPES, 5 mM MgSO_4_, 1 mM EGTA, 8 mg/ml BSA, pH 7.0), 10 μM biotin, 100 μM ATP, 100 μM THP, 2 μM Paclitaxel, and an oxygen scavenging system^45,46^ (final concentrations in buffer: 8 mg/mL glucose, 0.15 mg/mL catalase (from *Aspergillus niger*: Millipore Sigma, formerly EMD Millipore, 219261-100KU, 5100 U/mg), 0.29 mg/mL pyranose oxidase (from *Coriolus **sp.***: Sigma P4234-250UN, 12.2 U/mg), 100 μM Tris-HCl and 100 μM NaCl). In this trapping channel, dual-trap optical tweezers were used to trap a d-ssDNA-coated streptavidin microsphere in one trap, and a microsphere (Spherotech) coated with anti-digoxigenin antibody (Roche Diagnostics) in the other. The microspheres were repeatedly brought together until a d-ssDNA tether formed between them.

Once a d-ssDNA tether formed between the two trapped microspheres, force-extension curves (FECs) were collected by moving one trap away from the other at a constant rate (10 nm/s or 100 nm/s) over a pre-set distance, then returning at the same rate to the initial position. Multiple FECs were collected per tether, preferably until the tether ruptured, allowing us to observe the variability of the behavior of a single molecule. We determined whether a tether had consisted of a single molecule from the single, one-step rupture of the tether. At the low d-ssDNA concentrations used, only a small fraction of bead pairs (approximately 10%) formed tethers, decreasing the probability of multiple tethers forming.

### Optical Trapping Data Processing

Before the FECs could be used for calibration of d-ssDNA as a force sensor, the extension measured in optical trapping experiments was offset-corrected. The measured extension differs slightly from the absolute end-to-end extension of the molecule, owing to the difference in nominal and actual microsphere diameter and an instrumental offset. For measurements of a molecule with well-characterized stretching behavior, this offset is determined and corrected through fitting its FEC to the appropriate theoretical model, with the extension offset as the single fitting parameter. In the absence of such a model for d-ssDNA, we determined the offset (under the same conditions) by fitting the FEC of a well-characterized DNA hairpin construct to the extensible worm-like chain model^47,48^. The average offset was determined to be 60 nm. Variations in microsphere diameter were small: the standard deviation of the extension offset was 5 nm for the control construct, and the control set of FECs spanned ~25 nm at 15 pN. All reported d-ssDNA extensions were offset-corrected by 60 nm.

Out of 72 collected FECs, 64 formed a cluster with similar curvature (Fig. S4). We interpreted this ‘primary cluster’ as representing the typical behavior of d-ssDNA. The vast majority of these, 58 FECs, had no significant rips or discontinuities; this set was used to determine a calibration force-extension curve for d-ssDNA for the kinesin experiments. First, a single ‘net’ FEC for each bead pair was obtained from a weighted average of all FECs, binned in force, from that bead pair. The resulting six net FECs were fitted to a model based on previously published data and models of d-ssDNA behavior^28,31^. These published data show that the extension *L* of d-ssDNA under force *f* follows a power law, *L* ~ *f^γ^* with *γ* ≈ 2/3 at low forces, in solutions containing either monovalent or divalent ions^28,31^. At higher forces, *L* varies logarithmically with force in solutions containing monovalent ions only. Inspection of the rescaled FECs in solutions containing divalent ions (Fig. 3 of Ref.^31^) suggests that the extension may follow a power law with a different exponent over at least part of the higher force range. We therefore fitted our optical trapping data to the following expression:

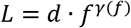

in which the force-dependent exponent *γ* transitions between two values: the experimentally determined value 0.62 at low force^31^mc, and the value 0.62 – *a* at high force:

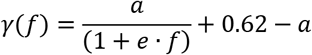

The FECs of d-ssDNA were fit to this expression with *a, d*, and *e* as fitting parameters. The resulting coefficients, with standard errors, are: *a* = 0.365 ± 0.003, *d* = 273 ± 2, and *e* = 5.6 ± 1.5.

Fitting was carried out on six averaged FECs, one per bead pair—rather than directly on the set of 58 FECs—to avoid biasing the fits through properties of beads that yielded more FECs. Factors specific to each bead pair, in particular any deviation of the beads’ diameters from the average values, and the resulting errors in calibration of the optical trap stiffness, affect all FECs from that bead pair in the same manner; these FECs cannot therefore strictly be treated as independent. Instead, each FEC from a given bead pair was binned by force; the mean extension and standard error of the mean extension in each bin was used to calculate a weighted average of extension from all FECs from the bead pair for that bin. The resulting extension per force bin comprised the net FEC for the bead (Fig. 2c, colored symbols; uncertainties of weighted averages are smaller than marker size). The resulting six net FECs were then fitted with the model with an unweighted fit. Force-extension data below 0.2 pN were not included in the fit: at those forces, the molecular extension for this construct is sufficiently low that the measurements become inaccurate due to artefacts, in particular cross-talk between the two optical traps. Our obtained analytical model for f is a transcendental equation with no closed from analytical solution for f, and, therefore a polynomial was fitted to our expression and was used to calculate the forces from displacements.

### Stretching behavior of d-ssDNA: variability and clustering

Figure S4 shows all FECs collected, color-coded by bead pair (10 in total). While tethers from a given bead pair may be different molecules, some may be previously measured molecules re-tethered. Out of the 72 collected FECs of d-ssDNA, 64 fell into a ‘primary’ cluster, which we interpreted as the typical stretching behavior of the molecule.

The primary cluster exhibits considerable variation, resulting in a range of extensions at any given force. Some of this variability may arise from differences in microsphere diameter and from the resulting errors in calibration of the optical trap stiffness. However, we see appreciable variability in FECs taken from the same bead pair—pointing to heterogeneity in d-ssDNA molecules—and sometimes even from the same molecule pulled through multiple cycles. The variability in FECs can be considerable, spanning up to 40 or 50 nm in extension at 15 pN. We see no evidence of significant secondary structure formation, which could account for the variability. Variability in molecular stretching behavior of d-ssDNA has been observed previously. McIntosh and Saleh report variations in contour length from molecule to molecule and also between FECs of the same molecule, especially at high salt concentrations^31^. These variations were attributed to non-specific adsorption to surfaces, but it is unclear if this would be relevant in our optical trap setup.

The remaining FECs fall into a ‘secondary’ cluster, with a higher stiffness than those in the primary cluster, and were excluded from the analysis. The nature of the secondary cluster is unclear, but it is unlikely that most of it results from multiple tethers pulled in parallel. In order to maximize the probability of stretching unique molecules, very low concentrations of d-ssDNA were used, and only ~10% of the microsphere pairs formed tethers. Moreover, some secondary FECs ruptured in one step. We suspect that most secondary FECs could arise from rare intra- or inter-molecular cross-linking events during synthesis^31,49^, or from the variability in contour length described above.

### Force sensor assay

Mammalian biotinylated kinesin (888 aa long) was purified from mouse cells and was a generous gift from Prof. Kathleen Trybus (U Vermont). Kinesin was mixed with 5x more quantum dots (QD705 streptavidin conjugate, Catalog Number: Q10163MP, ThermoFisher). Kinesin conjugated with quantum dots (Kin-QD) was purified from the free QDs using the in house developed MiCA purification^27^. Kin-QD was mixed with d-ssDNA in 5x molar excess to maximize the probability of attachment of one d-ssDNA molecule per kin-QD molecule (kin-QD-dssDNA). The biotin end of d-ssDNA got attached to kin-QD, and the digoxigenin end remained free. Kin-QD-dssDNA was mixed with 1 mM biotin to saturate all streptavidin on the surface of the QD.

Twenty-two square-mm coverslips were covalently functionalized with PEG and 1% PEG-Biotin^50^. Flow chambers were made by sandwiching double-sided tape between cleaned glass slides and PEG-functionalized coverslips. 600 nM streptavidin (Cat. No: 43-4302 from Thermo Scientific) was flowed in the flow chamber followed by wash by DMB-BSA (DMB buffer: 30 mM HEPES, 5 mM MgSO_4_, 1 mM EGTA, 8 mg/ml BSA, pH 7.0). 10 nM of anti-digoxigenin-biotin was flowed in the chamber and washed after 5 minutes with DMB-BSA-biotin buffer (DMB-BSA buffer supplemented with 0.5 mM biotin). Kin-QD-dssDNA was flowed in the chamber so that the assembly attaches the surface with the linkage of digoxigenin on d-ssDNA and anti-digoxigenin on the surface. Kin-QD-d-ssDNA solution was diluted sufficiently such that kinesins on the coverslips can be tracked individually and do not fall within the diffraction limit distance. After the attachment of kin-QD-d-ssDNA on the surface, imaging buffer containing microtubules, 2 mM ATP, and deoxygenating agents (pyranose oxidase+glucose) was flowed into the chamber (final concentrations in buffer: 8 mg/mL glucose, 0.15 mg/mL catalase (from *Aspergillus niger*: Millipore Sigma, formerly EMD Millipore, 219261-100KU, 5100 U/mg), 0.29 mg/mL pyranose oxidase (from *Coriolus **sp***.: Sigma P4234-250UN, 12.2 U/mg)). The flow chamber was immediately imaged after flowing of the imaging buffer. 1 μl of 50 nM of QD605 streptavidin conjugate (Catalog Number: Q10103MP, ThermoFisher) was mixed with 1 μl of biotinylated microtubules (~6 μM of microtubules) for mimicking the roadblocks. The qd605-microtubule mixture was incubated for 20 minutes. Streptavidin molecules on the surface of QD605 were saturated with 1 mM biotin. QD605 coated microtubules were used for making the imaging buffer. Force sensor assay was performed as described before with imaging buffer that contained roadblock incubated microtubules. For the experiments involving 100 nm roadblocks, we used 100 nm streptavidin-coated fluorescent polystyrene particles (Kisker Biotech, Germany). We mixed 3x diluted nanoparticles with 1 μl of microtubules (~6 μM of microtubules) and incubated for 10 mins. Microtubule-particle assembly was saturated with 1 mM biotin and was used in the experiment. For tau experiments, we mixed Alexa532 conjugated 3RS tau with the imaging buffer for the final concentration of 100 nm tau. We incubated the imaging buffer for 20 minutes before imaging it under the microscope.

### Fluorescence Imaging

Kinesin was labeled with QD705, and microtubules were labeled with hilyte488 and were simultaneously imaged on two camera system in total internal refraction fluorescence (TIRF) mode. Experiments were performed on an inverted light microscope (Olympus IX71) with Andor EMCCD (iXon DU-897E) cameras. Images were acquired at 100x magnification with oil immersion objective (Olympus UPlanSApo, NA 1.40). The sample was excited by a 488 nm blue laser (Coherent OBIS), and excitation light was reflected with 495 nm long-pass dichroic (Chroma). MultiCam (Cairn Research, UK) was used to split the kinesin and microtubule excitation signals to capture on two different cameras. The excitation signal was passed through a 685 nm (T685lpxr-UF3, UltraFlat, Chroma) long-pass filter to split kinesin and microtubule signals. Additional filters were installed for kinesin (710/40, Brightline, Semrock) as well as microtubule (560/80, Brightline, Semrock) signals just before the camera captured the image. Alexa532 labeled tau was excited by green laser (Coherent OBIS). Two thousand frames per movie were acquired at 0.2 second exposure time and variable EM gain. EM gain was chosen to get the maximal signal output without saturating the camera on a case by case basis.

For doing the roadblock experiment, three cameras on MultiCam were used to image kinesin, roadblocks, and microtubules simultaneously. QD605 were attached to microtubules using streptavidin-biotin linkage that mimicked roadblocks. The excitation signal was split first with a 670 nm long-pass filter. Fluorescence corresponding to wavelengths >685 was captured on the kinesin camera. Excitation light with wavelengths <685 nm was further split using 570 nm long pass filter for roadblock and microtubule channel. Light entering the roadblock channel was cleaned with an additional 600/80 (Brightline, Semrock) filter.

### Analysis of acquired movies

Movies obtained from different cameras were registered using nanohole and in-house MATLAB programs. Kinesin displacements and microtubule velocity were also calculated using the TrackMate^51^ plugin of ImageJ^52^ and in-house MATLAB, as described previously^6^. Force applied by individual kinesin was calculated using a polynomial fit of the model described in Fig. 2c with a MATLAB code.

## Data availability

Data is available in the main text or the supplementary information text and files. All Single-molecule imaging raw datasets are available from the corresponding author upon request.

## Materials availability

Reagents described in this work will be made available upon request to the authors.

## Code availability

The custom codes for the data analysis used in this study are available from the corresponding author upon request.

## Acknowledgments

This work was supported by the National Institute of Health grants R01 GM132392 (to P.R.S.), GM132646 (to C.L.B.), GM120353 (to Y.R.C.), R35 GM136288 (to K.M.T.) and by the National Science Foundation grant #1430124.

## Author contributions

Conceptualization: S.S., Y.R.C., and P.R.S.; methodology: S.S., A.T., Y.R.C., and P.R.S.; Single-molecule imaging experiments: S.S., N.S., and B.K.M.; d-ssDNA synthesis experiments: S.S.; Optical trap experiments and model fitting: A.T and Y.R.C.; Kinesin purification: C.S.B. and K.T.; Tau purification and labeling: L.C. and C.B.; MATLAB scripts: S.S. and M.T.; Imaging data analysis: S.S., N.S., and B.K.M.; Writing: S.S., A.T. Y.R.C., and P.R.S.; supervision: Y.R.C. and P.R.S.

## Supplementary information

Supplementary information

Supplementary movie 1-4

## References

1. Schliwa, M. & Woehlke, G. Molecular motors. Nature 422, 759–765 (2003).

2. Howard, J. Molecular motors: Structural adaptations to cellular functions. Nature 389, 561–567 (1997).

3. Habermann, A., Schroer, T. A., Griffiths, G. & Burkhardt, J. K. Immunolocalization of cytoplasmic dynein and dynactin subunits incultured macrophages: Enrichment on early endocytic organelles. J. Cell Sci. 114, 229–240 (2001).

4. Leopold, P. L., McDowall, A. W., Pfister, K. K., Bloom, G. S. & Brady, S. T. Association of kinesin with characterized membrane-bounded organelles. Cell Motil. Cytoskeleton 23, 19–33 (1992).

5. Gross, S. P., Vershinin, M. & Shubeita, G. T. Cargo Transport: Two Motors Are Sometimes Better Than One. Curr. Biol. 17, 478–486 (2007).

6. Tjioe, M. et al. Multiple kinesins induce tension for smooth cargo transport. Elife 8, 1–31 (2019).

7. Siahaan, V. et al. Kinetically distinct phases of tau on microtubules regulate kinesin motors and severing enzymes. Nat. Cell Biol. 21, 1086–1092 (2019).

8. Chaudhary, A. R., Berger, F., Berger, C. L. & Hendricks, A. G. Tau directs intracellular trafficking by regulating the forces exerted by kinesin and dynein teams. Traffic 19, 111–121 (2018).

9. Ferro, L. S., Can, S., Turner, M. A., Elshenawy, M. M. & Yildiz, A. Kinesin and dynein use distinct mechanisms to bypass obstacles. Elife (2019). doi:10.7554/eLife.48629

10. Svoboda, K. & Block, S. M. Force and velocity measured for single kinesin molecules. Cell 77, 773–784 (1994).

11. Coppin, C. M., Pierce, D. W., Hsu, L. & Vale, R. D. The load dependence of kinesin’s mechanical cycle. Proc. Natl. Acad. Sci. 94, 8539–8544 (2002).

12. Kunwar, A., Vershinin, M., Xu, J. & Gross, S. P. Stepping, Strain Gating, and an Unexpected Force-Velocity Curve for Multiple-Motor-Based Transport. Curr. Biol. 18, 1173–1183 (2008).

13. Blehm, B. H., Schroer, T. A., Trybus, K. M., Chemla, Y. R. & Selvin, P. R. In vivo optical trapping indicates kinesin’s stall force is reduced by dynein during intracellular transport. Proc. Natl. Acad. Sci. 110, 3381–3386 (2013).

14. Rai, A. K., Rai, A., Ramaiya, A. J., Jha, R. & Mallik, R. Molecular adaptations allow dynein to generate large collective forces inside cells. Cell 152, 172–182 (2013).

15. Kuo, S. C. & Sheetz, M. P. Force of single kinesin molecules measured with optical tweezers. Science 260, 232–234 (1993).

16. Furuta, K., Furuta, A. & Toyoshima, Y. Y. Measuring collective transport by defined numbers of processive and nonprocessive kinesin motors. Proc. Natl. Acad. Sci. 110, 501–506 (2013).

17. Blanchard, A. T. & Salaita, K. Emerging uses of DNA mechanical devices. Science 365, 1080–1081 (2019).

18. Albrecht, C. et al. DNA: A programmable force sensor. Science 301, 367–370 (2003).

19. Wang, X. & Ha, T. Defining single molecular forces required to activate integrin and Notch signaling. Science 340, 991–994 (2013).

20. Brockman, J. M. et al. Mapping the 3D orientation of piconewton integrin traction forces. Nat. Methods 15, 115–118 (2018).

21. Lilley, D. M. J. & Ha, T. Fluorescence-Force Spectroscopy Maps Two-Dimensional Reaction Landscape of the Holliday Junction. Science 1909, 279–284 (2007).

22. Funke, J. J. et al. Uncovering the forces between nucleosomes using DNA origami. Sci. Adv. 2, e1600974 (2016).

23. Nickels, P. C. et al. Molecular force spectroscopy with a DNA origami-based nanoscopic force clamp. Science 354, 305–307 (2016).

24. A programmable DNA origami nanospring that reveals force-induced adjacent binding of myosin VI heads Iwaki, M., Wickham, S. F., Ikezaki, K., Yanagida, T. & Shih, W. M. A programmable DNA origami nanospring that reveals force-induced adjacent binding of myosin VI heads. Nat. Commun. 7, 1–10 (2016).

25. Yildiz, A. et al. Myosin V Walks Hand-Over-Hand: Single Fluorophore Imaging with 1.5-nm Localization. Science 300, 2061–2065 (2003).

26. McMaster, G. K., Carmichael, G. G. & Gordon, G. Analysis of single- and doublestranded nucleic acids on polyacrylamide and agarose gels by using glyoxal and acridine orange. Proc. Natl. Acad. Sci. U. S. A. 74, 4835–4838 (1977).

27. Tjioe, M. et al. Magnetic Cytoskeleton Affinity (MiCA) Purification of Microtubule Motors conjugated to Quantum Dots. Bioconjug. Chem. 29, 2278–2286 (2018).

28. Saleh, O. A., McIntosh, D. B., Pincus, P. & Ribeck, N. Nonlinear low-force elasticity of single-stranded DNA molecules. Phys. Rev. Lett. 102, 1–4 (2009).

29. Moffitt, J. R., Chemla, Y. R., Izhaky, D. & Bustamante, C. Differential detection of dual traps improves the spatial resolution of optical tweezers. Proc. Natl. Acad. Sci. U. S. A. 103, 9006–11 (2006).

30. Bustamante, C., Chemla, Y. R. & Moffitt, J. R. High-resolution dual-trap optical tweezers with differential detection: Instrument design. Cold Spring Harb. Protoc. 4, (2009).

31. McIntosh, D. B. & Saleh, O. A. Salt species-dependent electrostatic effects on ssDNA elasticity. Macromolecules 44, 2328–2333 (2011).

32. Hunt, A. J., Gittes, F. & Howard, J. The force exerted by a single kinesin molecule against a viscous load. Biophys. J. 67, 766–781 (1994).

33. Schnitzer, M. J., Visscher, K. & Block, S. M. Single kinesin molecules studied with a molecular force clamp. Nature 400, 184–189 (1999).

34. Nishiyama, M., Higuchi, H. & Yanagida, T. Chemomechanical coupling of the forward and backward steps of single kinesin molecules. Nat. Cell Biol. 4, 790–797 (2002).

35. Belyy, V. et al. The mammalian dynein–dynactin complex is a strong opponent to kinesin in a tug-of-war competition. Nat. Cell Biol. (2016). doi:10.1038/ncb3393

36. Albe, K. R., Butler, M. H. & Wright, B. E. Cellular concentrations of enzymes and their substrates. J. Theor. Biol. 143, 163–195 (1990).

37. Berret, J.-F. Local viscoelasticity of living cells measured by rotational magnetic spectroscopy. Nat. Commun. 7, 10134 (2016).

38. Wang, Y. & Mandelkow, E. Tau in physiology and pathology. Nat. Rev. Neurosci. 17, 5–21 (2016).

39. Stern, J. L., Lessard, D. V., Hoeprich, G. J., Morfini, G. A. & Berger, C. L. Phosphoregulation of Tau modulates inhibition of kinesin-1 motility. Mol. Biol. Cell 28, 1079–1087 (2017).

40. Tan, R. et al. Microtubules gate tau condensation to spatially regulate microtubule functions. Nat. Cell Biol. 21, 1078–1085 (2019).

41. Monroy, B. Y. et al. Competition between microtubule-associated proteins directs motor transport. Nat. Commun. 9, 1–12 (2018).

42. Ferro, L. S. et al. The mechanism of motor inhibition by microtubule-associated proteins. bioRxiv 2020.10.22.351346 (2020). doi:10.1101/2020.10.22.351346

43. Dixit, R., Ross, J. L., Goldman, Y. E. & Holzbaur, E. L. F. Differential Regulation of Dynein and Kinesin Motor Proteins by Tau. Science 319, 1086–1089 (2008).

44. Whitley, K. D., Comstock, M. J. & Chemla, Y. R. High-Resolution Optical Tweezers Combined With Single-Molecule Confocal Microscopy. Methods in Enzymology 582, (Elsevier Inc., 2017).

45. Landry, M. P., McCall, P. M., Qi, Z. & Chemla, Y. R. Characterization of photoactivated singlet oxygen damage in single-molecule optical trap experiments. Biophys. J. 97, 2128–2136 (2009).

46. Swoboda, M. et al. Enzymatic oxygen scavenging for photostability without ph drop in single-molecule experiments. ACS Nano 6, 6364–6369 (2012).

47. Wang, M. D., Yin, H., Landick, R., Gelles, J. & Block, S. M. Stretching DNA with optical tweezers. Biophys. J. 72, 1335–1346 (1997).

48. Camunas-Soler, J., Ribezzi-Crivellari, M. & Ritort, F. Elastic Properties of Nucleic Acids by Single-Molecule Force Spectroscopy. Annu. Rev. Biophys. 45, 65–84 (2016).

49. Kasai, H., Iwamoto-Tanaka, N. & Fukada, S. DNA modifications by the mutagen glyoxal: Adduction to G and C, deamination of C and GC and GA cross-linking. Carcinogenesis 19, 1459–1465 (1998).

50. Roy, R., Hohng, S. & Ha, T. A practical guide to single-molecule FRET. Nature Methods 5, 507–516 (2008).

51. Tinevez, J. Y. et al. TrackMate: An open and extensible platform for single-particle tracking. Methods 115, 80–90 (2017).

52. Schindelin, J. et al. Fiji: An open-source platform for biological-image analysis. Nature Methods 9, 676–682 (2012).

