## Supplementary Information pdf for "Individual Molecular Motors use Low Forces to bypass Roadblocks during Collective Cargo Transport"

### for

#### Supplementary Text

##### S1: Effect of vertical distance of microtubule on the d-ssDNA extension

In the simple microtubule gliding assay (without a d-ssDNA linker and with, kinesin directly immobilized on the surface), the distance between the microtubule and the surface was measured to be ~ 17 nm[1]. The radius of gyration (Rg) represents the radius of any polymer chain in its equilibrium position. The vertical length of the d-ssDNA molecule will be ~2*Rg.

Rg for ssDNA is given by[2]:

$$R_g = \left(\frac{L.p}{3}\right)^{1/2}$$

Where L is the contour length, and p is the persistence length of the molecule.

Rg of 1180 bp long d-ssDNA comes out to be ~21 nm. Therefore, the approximate distance of the microtubule from the surface in the FSIM assay would be ~38 nm (17+21 nm). 38 nm vertical distance will have minimal effect on large kinesin displacements (and hence forces) in our assay. For small kinesin displacements, the force exerted by the kinesins will be relatively small (<1 pN), and therefore, the vertical distance will not alter forces significantly. Therefore, we have assumed that the displacement of kinesin is approximately equal to the d-ssDNA extension.

#### S2: Drag force on a microtubule

Typical drag force values on the microtubules (MTs) are very low (0.01-0.1 pN). Drag force on microtubule can be calculated from the following equation-

$$F = C1.\eta.L.v$$

Where F is the drag force, C1 is the drag coefficient, η is viscosity, L is the length of the microtubule, and v is MT velocity[3].

As a test case: assuming viscosity of water, 10-micron length of MT, and 1 micron/sec velocity of MT: The drag forces comes out to be ~0.06 pN. Therefore, what we are observing agrees with the typical drag forces on the MT.

#### S3: Sensitivity and dynamic range of FSIM assay

A force sensor with a dynamic range of 0-10 pN is appropriate for the experiments of molecular motors as the stall forces of kinesin and dynein are below 7 pN. FSIM assay works well in this force range. Optical trap experiments with d-ssDNA molecule inform us that FSIM assay can measure forces up to 20 pN (Fig. 2c). We note that the standard deviation of the force-extension curve increases as we go above 10 pN, and it may therefore introduce uncertainty in the force calculation. We predict that the FSIM assay could be useful for other systems with higher force ranges but will need improvements in d-ssDNA preparation, purification, and theoretical modeling of the force-extension characteristics.

In the 0-7 pN force range (relevant for molecular motors), some uncertainty in the force calculation is introduced by the localization precision of the quantum dot-kinesin (i.e., the calculation of the d-ssDNA extension). In our acquired datasets, the localization uncertainty of kinesin-QD is ~14 nm (Fig. S15). It translates to the force error of <0.1 pN for forces below 1 pN. The error of force calculation due to localization uncertainty increases gradually in the higher force range with the error of ~0.5 pN in 4-5 pN force range. Further improvements in the localization precision of the kinesin-quantum dots can reduce the error of force calculation in the FSIM assay.

#### Supplementary figures

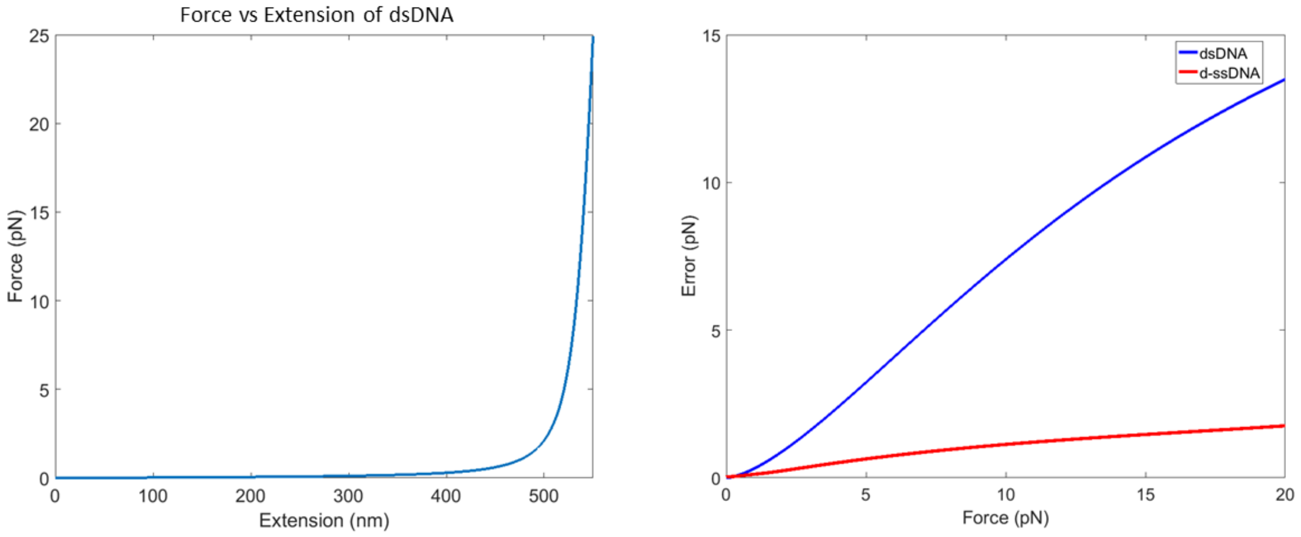


**Figure S1. dsDNA force-extension model and the error in force measurement.** The force required to stretch a 1.5 kb dsDNA molecule remains very small up to an extension of ~450 nm and shoots up near its contour length (left panel). After the extension of 450 nm, there is a large variation in force with a small change in extension (eWLC model)[4]. Given the average localization precision of ~14 nm (see fig. S15), the error in the calculation of the force is plotted for both dsDNA and d-ssDNA (right panel). dsDNA is not a reliable force sensing molecule because of the large errors in force calculation.

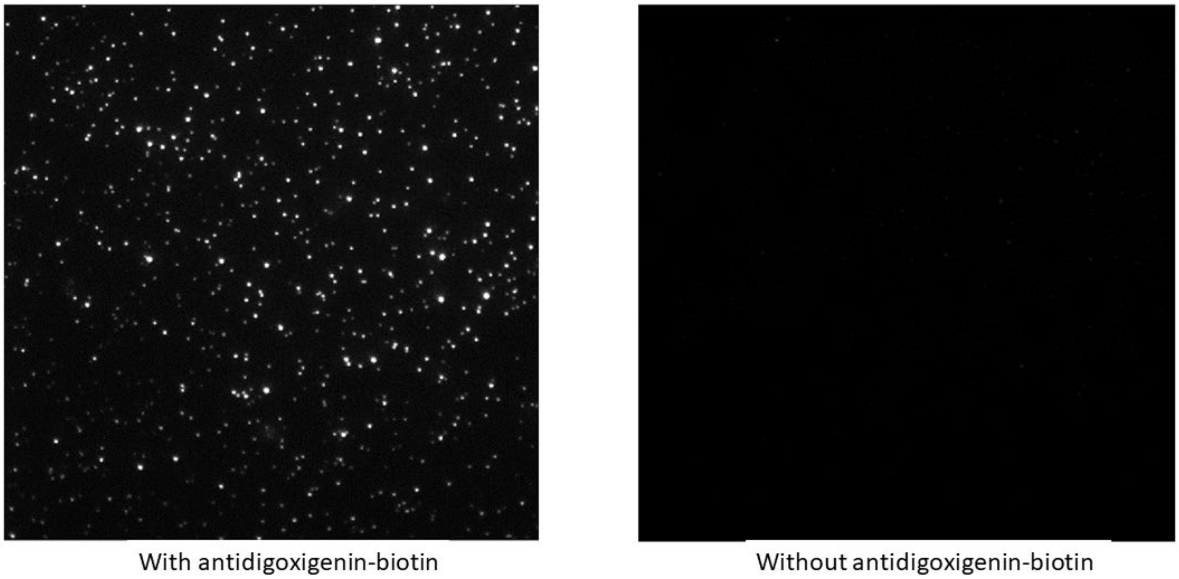


**Figure S2. Control for negligible non-specific binding of QDs**. The left image shows the concentration of kinesins on the surface when 1 nM of kinesin-QD is flown in the imaging chamber with anti-digoxigenin-biotin. Antidigoxigenin-biotin is immobilized on the surface using its biotin end, and the anti-digoxigenin end attaches to the digoxigenin end of the ssDNA-QD-kinesin. When kinesin-QD-ssDNA is flown in the chamber without anti-digoxigenin on the surface, ideally, there should be no kinesin attachment on the surface. This is what we observe in the right-side microscope image, proving there is the negligible non-specific binding of kinesin-QDs on the surface. In our force-sensing experiment, every kinesin has a d-ssDNA molecule attached to it.

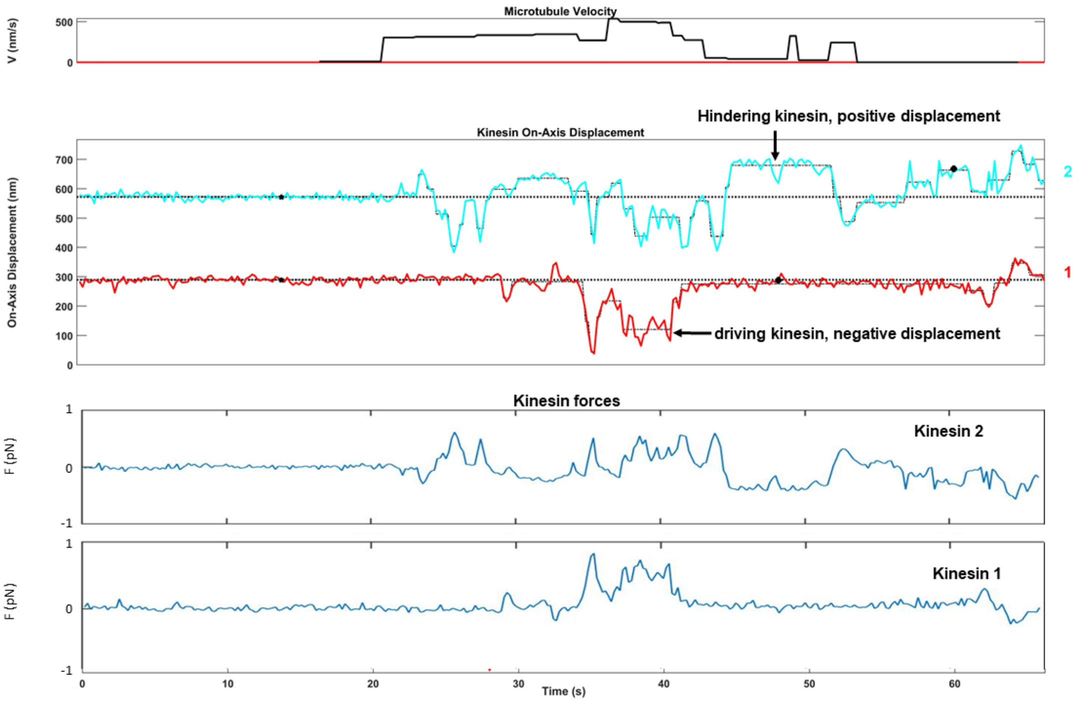


**Figure S3. Two kinesins transporting a microtubule.** Microtubule velocity, kinesin displacements, and kinesin forces are plotted with time. Kinesin can exhibit both negative and positive displacement and can adopt driving or hindering states. From kinesin displacement, we calculate kinesin forces. Kinesin can apply forces in both positive and negative directions.

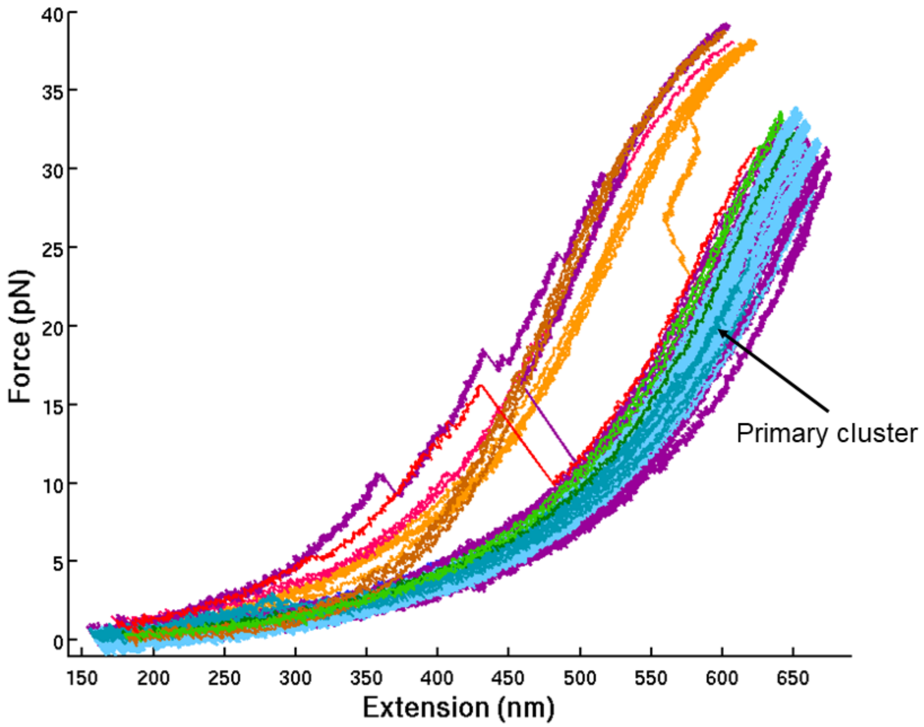


**Figure S4. Force extension curves of d-ssDNA.** All force-extension curves of d-ssDNA were obtained using an optical trap assay. FECs are colored by bead pairs. The majority of FECs make the "primary cluster". We used the primary cluster for modeling the force-extension analytical expression. Out of the 72 collected FECs of d-ssDNA, 64 fell into a 'primary' cluster, which we interpreted as the typical stretching behavior of the molecule.

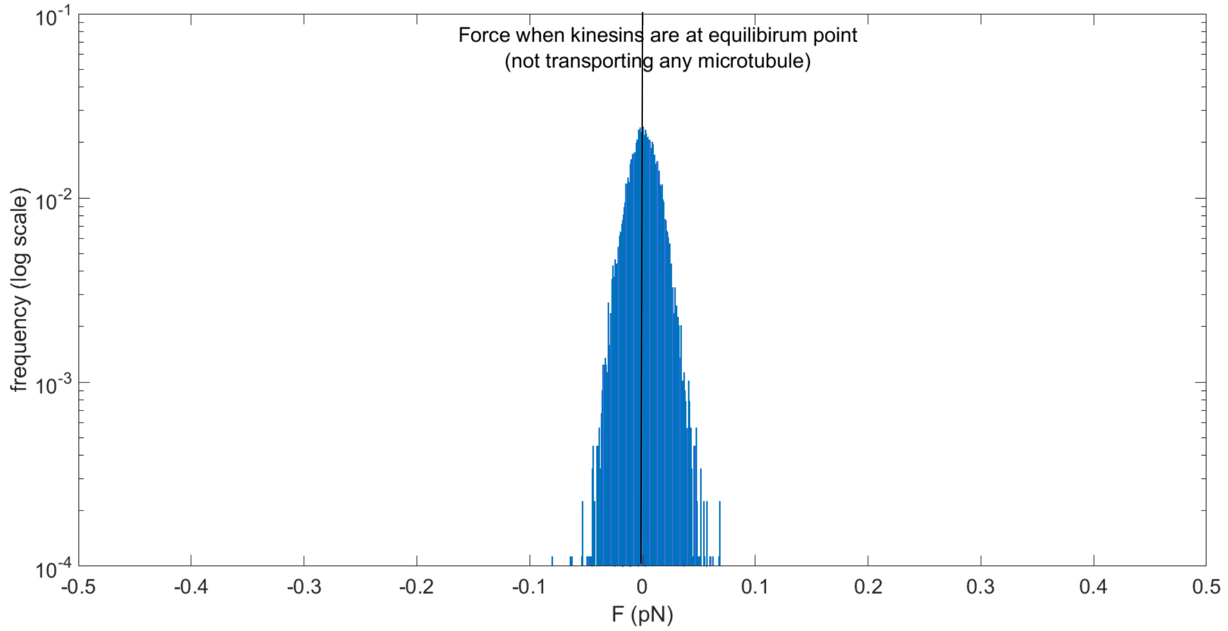


**Figure S5. Force by kinesins at the equilibrium position.** A large peak at zero forces is observed when there is no microtubule attached to the kinesin, and kinesins remain at their equilibrium positions.

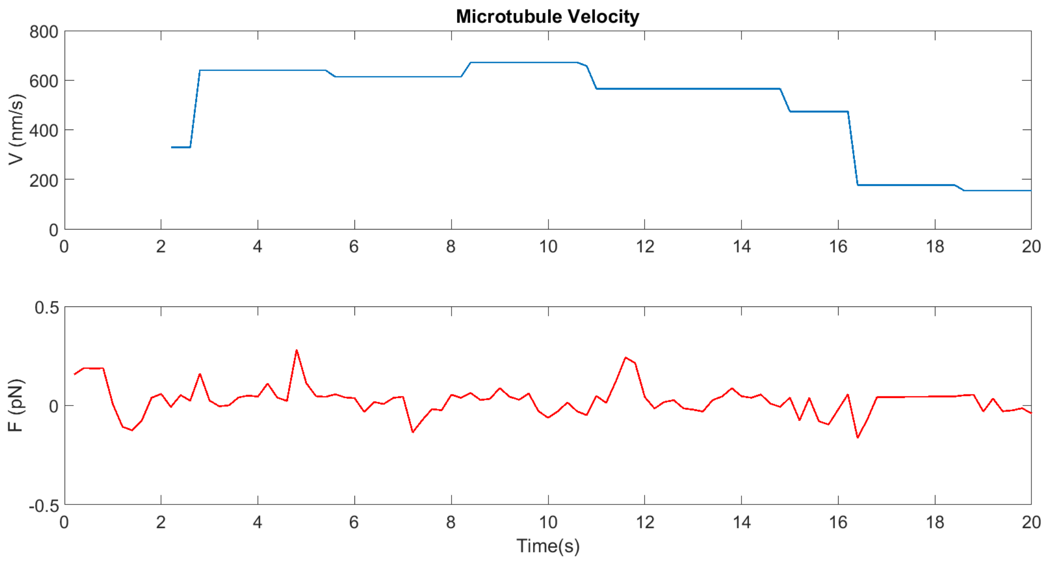


**Figure S6. Single kinesin transporting a microtubule.** Forces remain low when one kinesin transports a microtubule. Kinesin needs to act against the drag force on the microtubule, which is very low in the solution (see supplementary text 2).

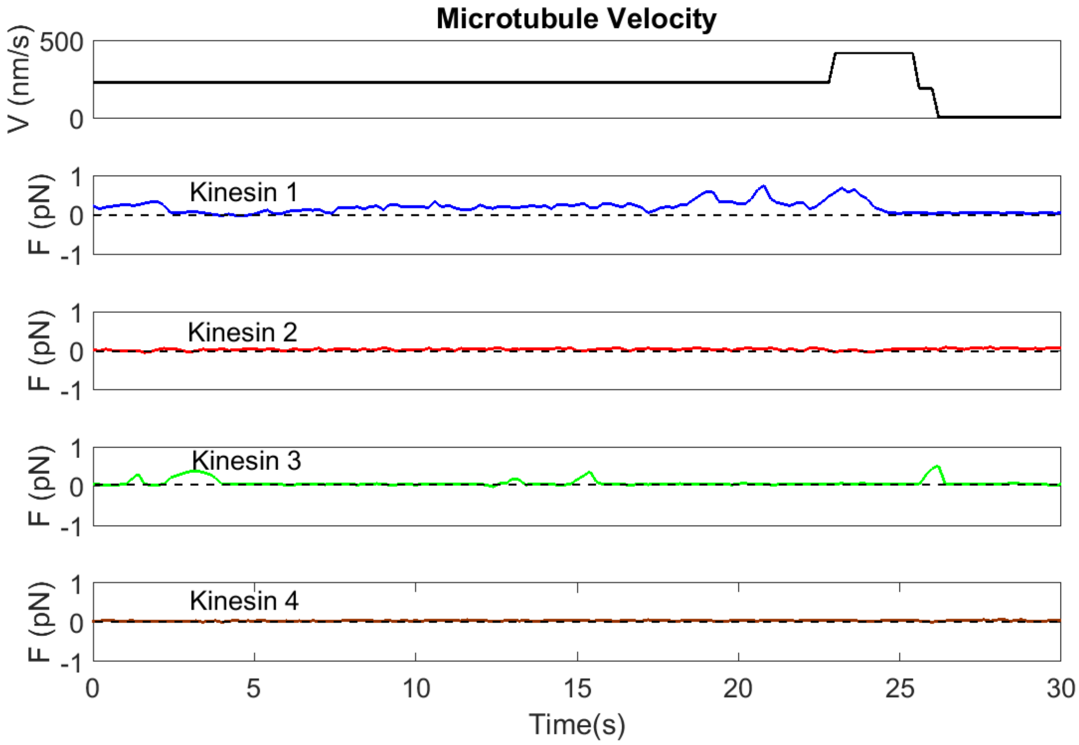


**Figure S7. Microtubule transported by four kinesins**. Individual forces by four kinesins along with the microtubule velocity are shown. Kinesin #1 and #3 occasionally exert positive forces, whereas kinesin #2 and #4 remain at the equilibrium position.

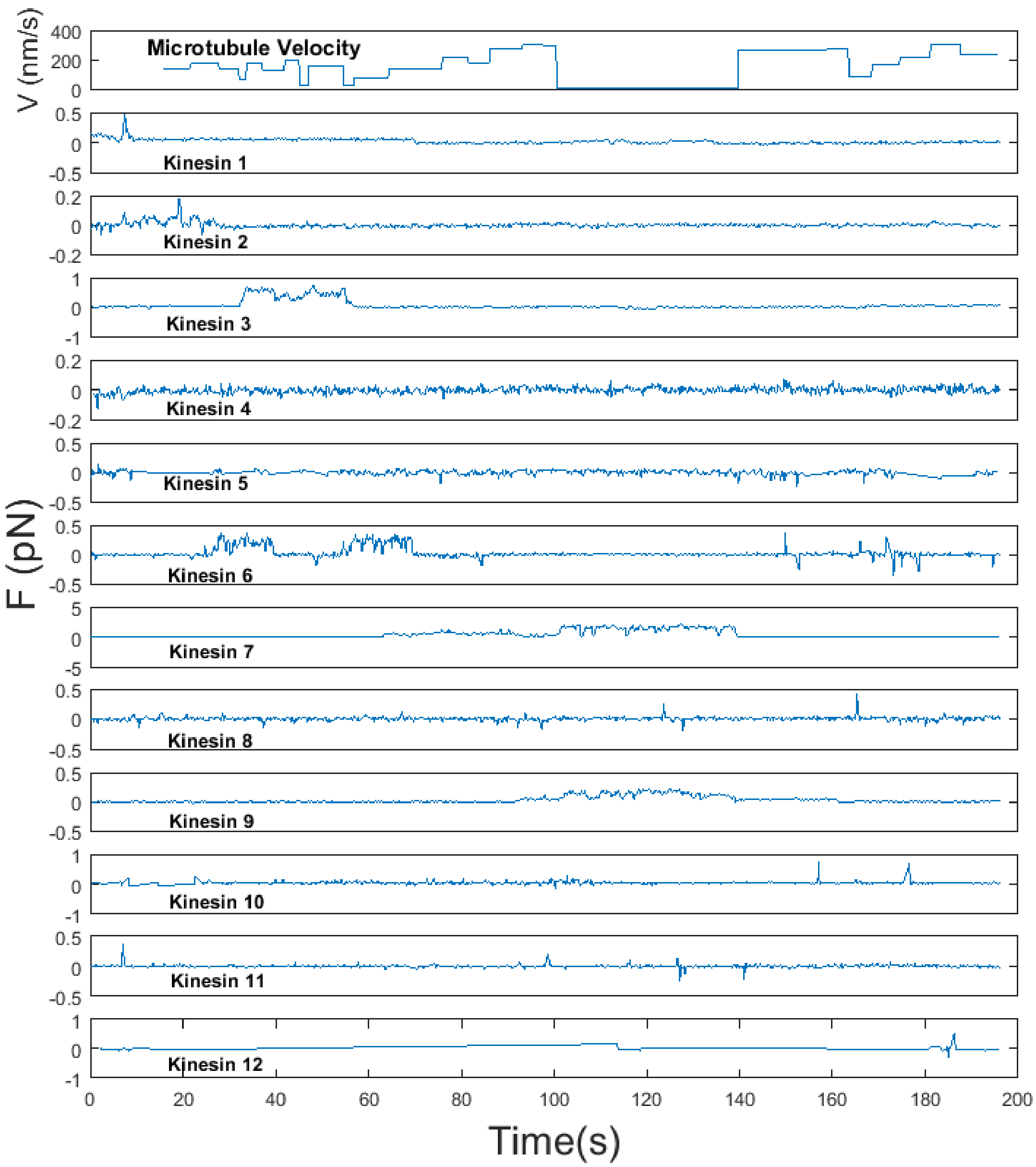


**Figure S8. Kinesin transported by 12 kinesins.** Forces of individual kinesins are plotted along with the microtubule velocity. Kinesin forces remain mostly below 1 pN during the cargo transport.

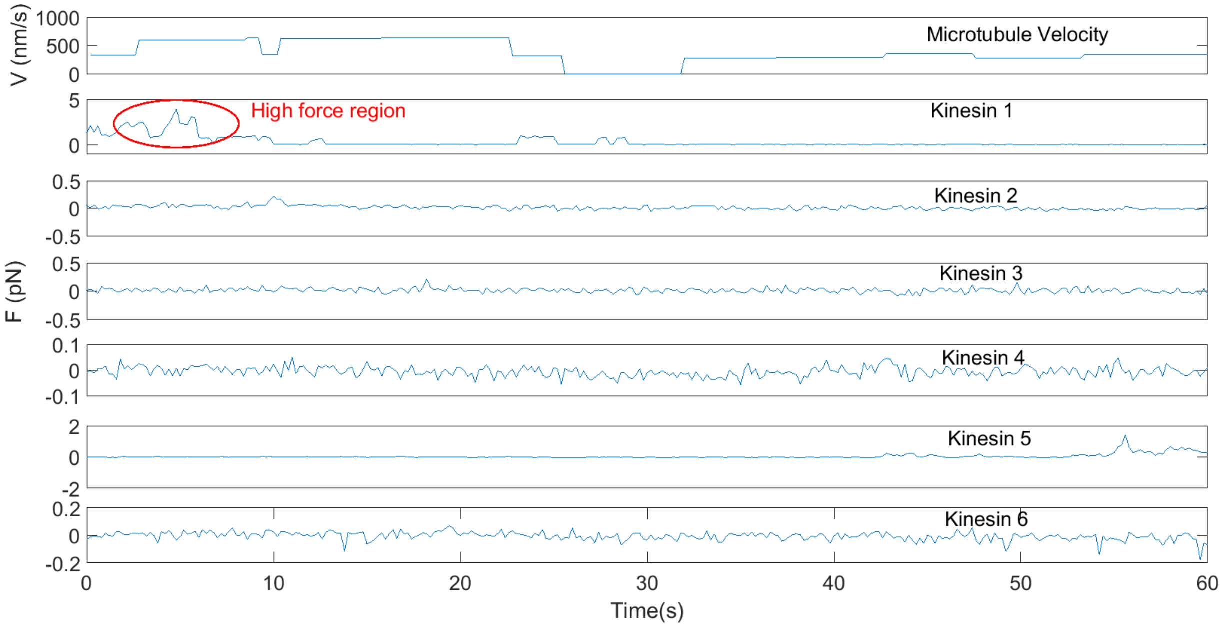


**Figure S9. Kinesin exerts more than a pN force.** A case where a microtubule is being transported by six kinesins (no roadblocks). Kinesin #1 produces ~4 pN of force at t=5 sec. This is one of the rare cases where a kinesin exerts more than one pN during cargo transport.

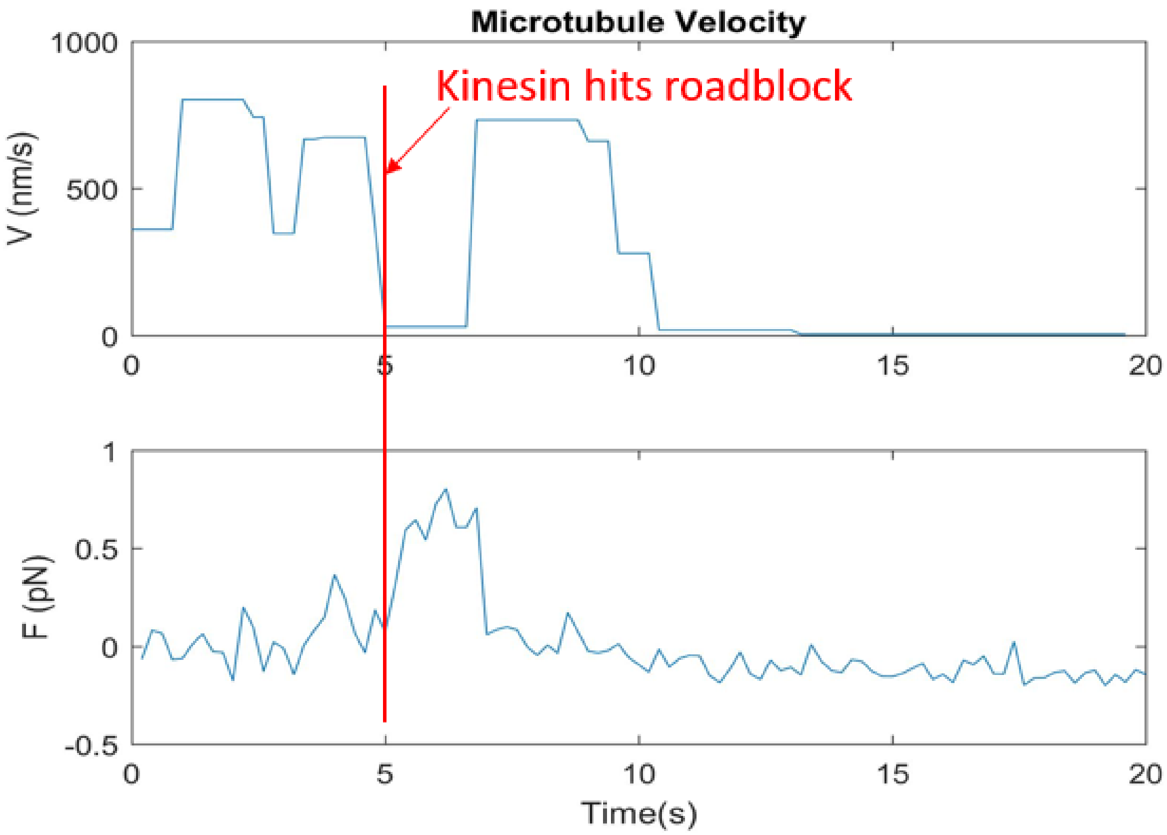


**Figure S10. One kinesin overcomes the roadblock.** A case where one kinesin is transporting the microtubule. Kinesin hits the roadblock (~20 nm quantum dot roadblock) at t=5 sec. It increases its force to overcome the roadblock.

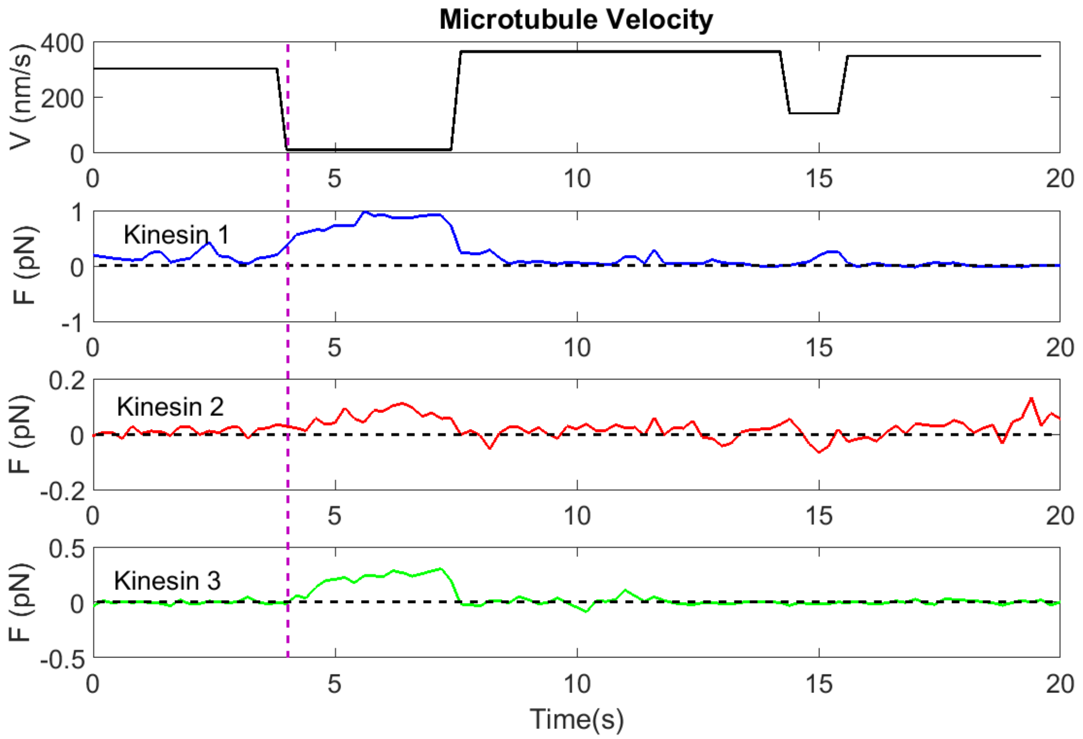


**Figure S11. Three kinesins resume microtubule motion.** In this example, the microtubule is stuck at the roadblock (~20 nm quantum dot roadblock) for a long time (from ~4-7 seconds). After t=4 sec, all three kinesins exert positive force on the microtubule, resuming its velocity at t=7 sec.

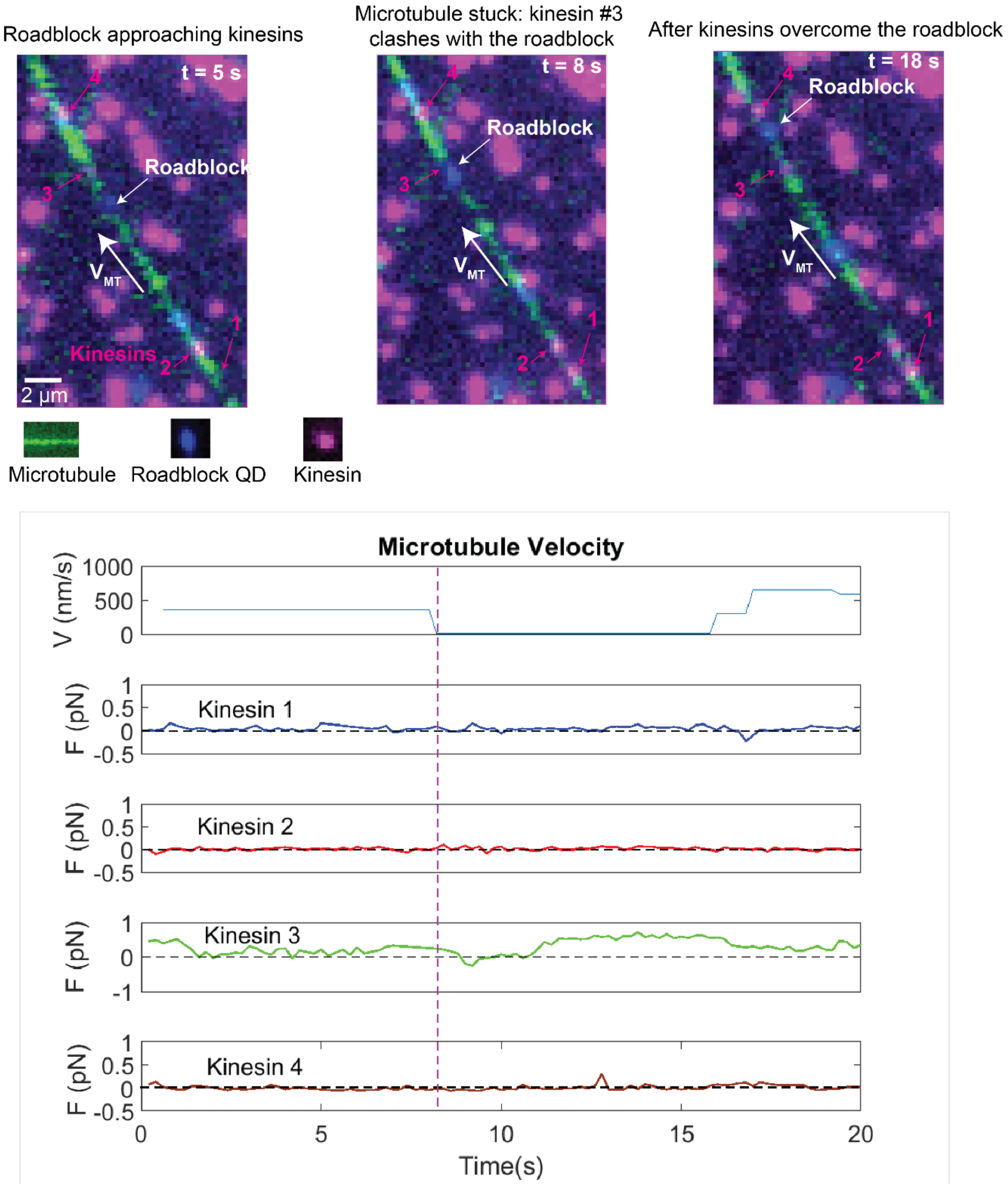


**Figure S12. Four kinesins overcome a roadblock.** Three microscope images (top) of an example are shown where a microtubule (shown in green) is transported by four kinesins (shown in magenta). Microtubule velocity and forces of individual kinesins are plotted (bottom). Kinesin#3 gets stuck at the roadblock (~20 nm quantum dot roadblock, shown in blue) for a long time (starting ~8 sec, vertical purple line). After t=16 sec, kinesin 3 exerts a positive force on the microtubule, resuming its velocity. The other three kinesins exert low forces.

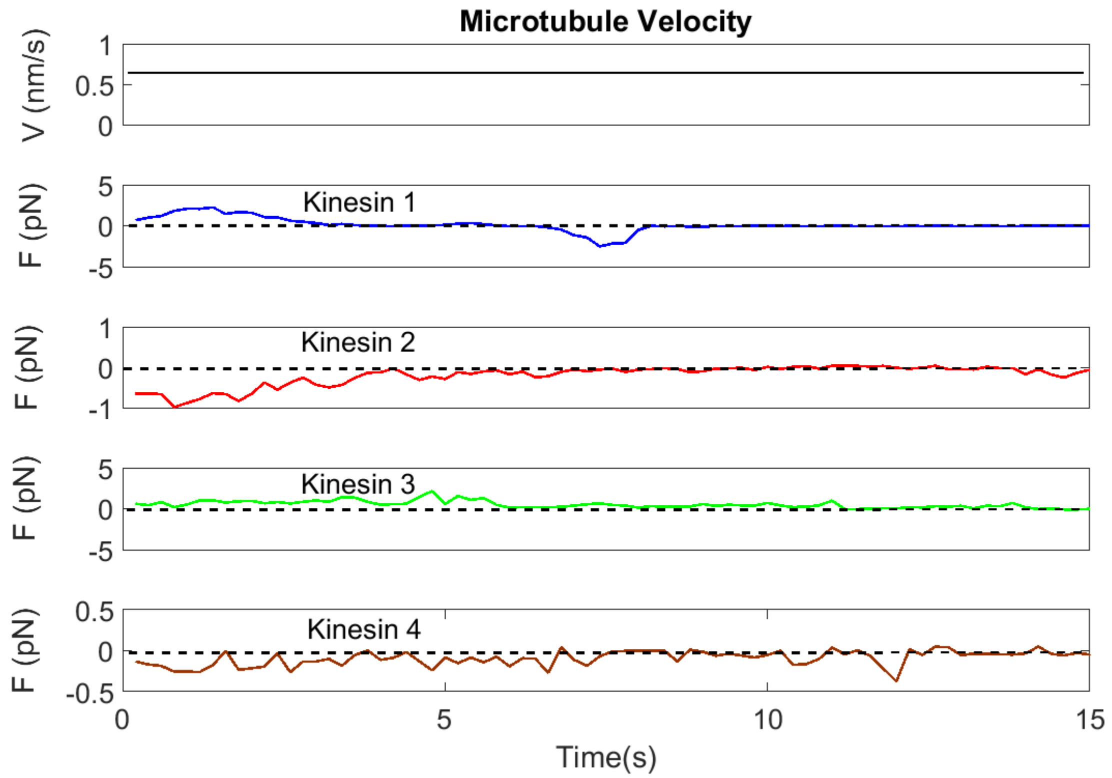


**Figure S13. Example of tau coated microtubule transport with four kinesins.** In this example, the tau coated microtubule transported by four kinesins and forces of individual kinesins is plotted with time.

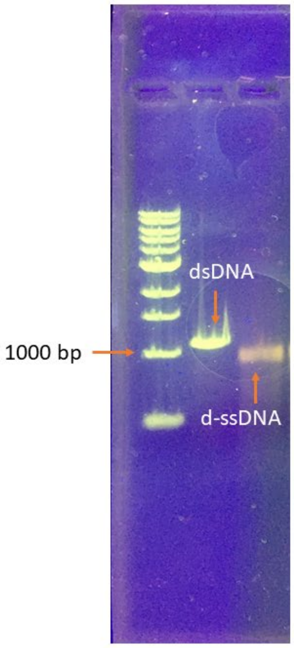


**Figure S14. Glyoxal denatured ssDNA (d-ssDNA).** The first lane is 1 kb dsDNA ladder. The second lane is control dsDNA (1080 bp). The third lane is d-ssDNA (1080 bp). The gel was stained with SYBR Gold dye.

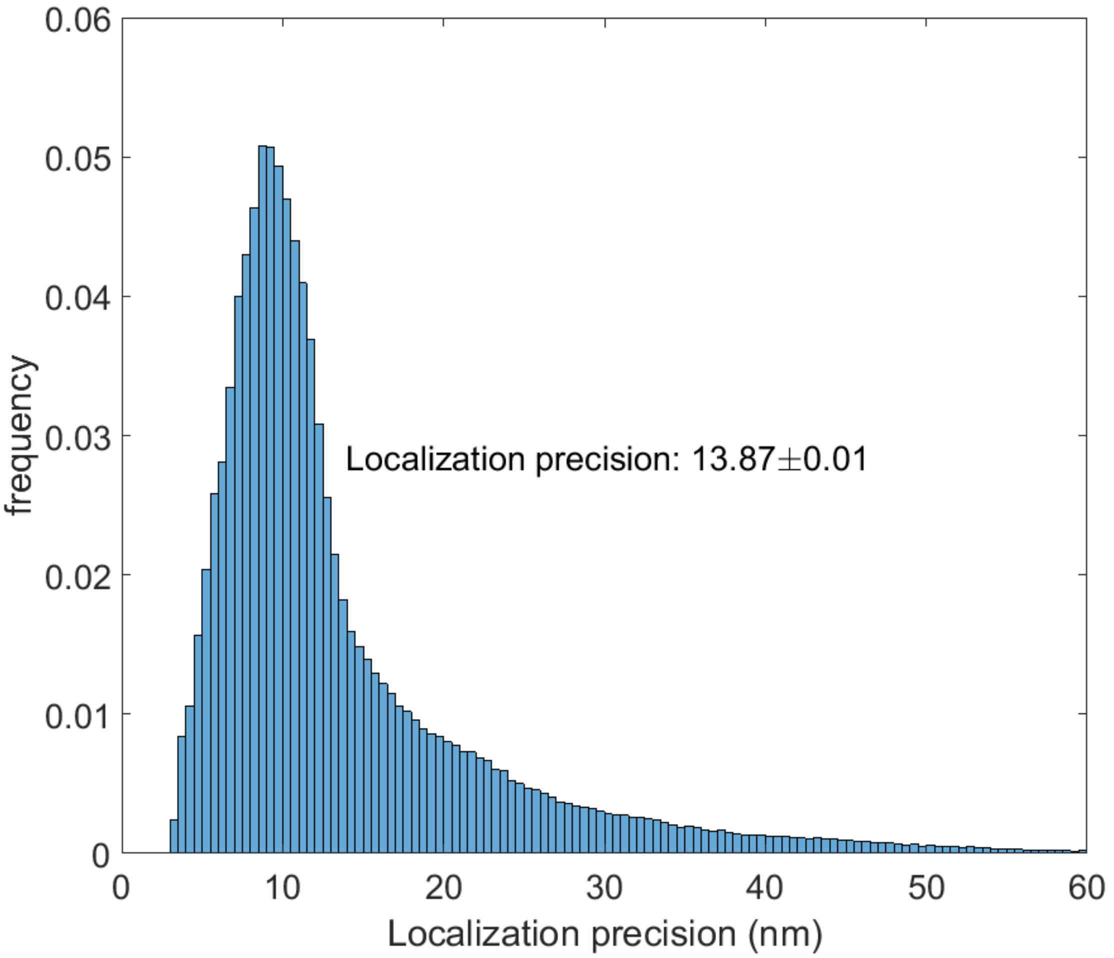


**Figure S15. Localization precision of kinesin-quantum dot**. The average localization precision of kinesin-QD is ~14 nm in our acquired data for FSIM assay **(**number of molecules analyzed=466,464) (Calculated by Thunderstorm Plugin[5] of Fiji[6]).

**SI References:**


1. Kerssemakers, J., Howard, J., Hess, H. & Diez, S. The distance that kinesin-1 holds its cargo from the microtubule surface measured by fluorescence interference contrast microscopy. *Proc. Natl. Acad. Sci. U. S. A.* **103**, 15812–15817 (2006).

2. Ochman, R. G. H. H. Persistence Length of Single-Stranded DNA. *Macromolecules* **30**, 5763–5765 (1997).

3. Hunt, A. J., Gittes, F. & Howard, J. The force exerted by a single kinesin molecule against a viscous load. *Biophys. J.* **67**, 766–781 (1994).

4. Lee, N.-K. & Thirumalai, D. Pulling-speed-dependent force-extension profiles for semiflexible chains. *Biophys. J.* **86**, 2641–9 (2004).

5. Ovesný, M., Křížek, P., Borkovec, J., Švindrych, Z. & Hagen, G. M. ThunderSTORM: A comprehensive ImageJ plug-in for PALM and STORM data analysis and super-resolution imaging. *Bioinformatics* **30**, 2389–2390 (2014).

6. Schindelin, J. *et al.* Fiji: An open-source platform for biological-image analysis. *Nature Methods* **9**, 676–682 (2012).
